# *Shigella flexneri* adherence factor expression in *in vivo*-like conditions

**DOI:** 10.1101/514679

**Authors:** Rachael B. Chanin, Kourtney P. Nickerson, Alejandro Llanos-Chea, Jeticia R. Sistrunk, David A. Rasko, Deepak Kumar Vijaya Kumar, John de la Parra, Jared R. Auclair, Jessica Ding, Kelvin Li, Snaha Krishna Dogiparthi, Benjamin J. D. Kusber, Christina S. Faherty

## Abstract

The *Shigella* species are Gram-negative, facultative intracellular pathogens that invade the colonic epithelium and cause significant diarrheal disease. Despite extensive research on the pathogen, comprehensive understanding of how *Shigella* initiates contact with epithelial cells remains unknown. *Shigella* maintains many of the same *Escherichia coli* adherence gene operons; however, at least one critical gene component in each operon is currently annotated as a pseudogene in reference genomes. These annotations, coupled with a lack of structures upon microscopic analysis following growth in laboratory media, have led the field to hypothesize that *Shigella* is unable to produce fimbriae or other “traditional” adherence factors. Nevertheless, our previous analyses have demonstrated that a combination of bile salts and glucose induce both biofilm formation and adherence to colonic epithelial cells. Through a two-part investigation, we first utilized various transcriptomic analyses to demonstrate that *S. flexneri* strain 2457T adherence gene operons are transcribed. Subsequently, we performed mutation, electron microscopy, biofilm, infection, and proteomic analyses to characterize three of the structural genes. In combination, these studies demonstrate that despite the gene annotations, *S. flexneri* 2457T uses adherence factors to initiate biofilm formation as well as epithelial cell contact. Furthermore, host factors, namely glucose and bile salts in the small intestine, offer key environmental stimuli required for proper adherence factor expression in *S. flexneri.* This research may have a significant impact on vaccine development for *Shigella* and further highlights the importance of utilizing *in vivo*-like conditions to study bacterial pathogenesis.

**Importance:** Bacterial pathogens have evolved to regulate virulence gene expression at critical points in the colonization and infection processes to successfully cause disease. The *Shigella* species infect the epithelial cells lining the colon to result in millions of cases of diarrhea and a significant global health burden. As antibiotic resistance rates increase, understanding the mechanisms of infection are vital to ensure successful vaccine development. Despite significant gains in our understanding of *Shigella* infection, it remains unknown how the bacteria initiate contact with the colonic epithelium. Most pathogens harbor multiple adherence factors to facilitate this process, but *Shigella* was thought to have lost the ability to produce these factors. Interestingly, we have identified conditions that mimic some features of gastrointestinal transit and enable *Shigella* to express adherence factors. This work highlights aspects of genetic regulation for *Shigella* adherence factors and may have a significant impact on future vaccine development.

## Introduction

*Shigella flexneri* is a Gram-negative, facultative anaerobe that infects millions of people each year by causing watery or bloody diarrhea, cramping, and dehydration. *Shigella* infection is endemic in developing countries, causing significant mortality and morbidity, particularly in children under the age of five years (1). In industrialized nations, infection is episodic and primarily linked to contaminated food or water. Infection in non-immunocompromised individuals is self-limiting and most patients recover with oral rehydration therapy and antibiotics (2–4). However, the increasing prevalence of antibiotic resistance (5) highlights the need to pursue effective vaccine strategies in these enteric pathogens that are gaining resistance mechanisms.

The current *Shigella* infection paradigm is that the bacteria spread through fecal-oral transmission in which an extremely low infectious dose, with as few as 10 to 100 organisms, initiates infection (2). Once ingested, *Shigella* traverses the digestive tract and localizes to the colon. To invade the colonic epithelium, *Shigella* transits through M (microfold or membranous) cells, which are specialized antigen-presenting cells of the follicle-associated epithelium (FAE) (6). Transit through M cells allows the bacteria to reach the basolateral pole of the epithelium for invasion (2), and the FAE is considered the major site of entry for *Shigella* due to the presence of M cells (7). Following basolateral invasion, intracellular replication, and intercellular spread, polymorphonuclear cells are recruited to the site of infection to eliminate the pathogen. The massive tissue destruction that results in the symptoms of bacillary dysentery is due to this intense inflammatory response (2).

While the invasion process and intracellular spread, replication, and survival of *Shigella* have been thoroughly investigated, much less is known about the virulence dynamics of the bacteria prior to invasion and transcytosis. In fact, there is a critical gap in knowledge regarding how the bacteria target M cells to initiate the invasion process and if *Shigella* utilizes adherence factors to adhere to the apical surface of epithelial cells prior to invasion. Due to the mucosal environment encountered on the surface of gastrointestinal epithelial cells, many pathogens, particularly pathogenic *Escherichia coli* and *Salmonella*, often utilize pili, fimbriae, or afimbrial adhesins to efficiently colonize host cells (8–13). Since *Shigella* evolved from *E. coli* (14, 15) and given the fact that fimbriae are prevalent among the *Enterobacteriaceae* (16), it is reasonable to hypothesize that *Shigella* utilize fimbriae or other adhesins during colonization. Interestingly, *Shigella* is thought to have lost the ability to produce “traditional” *E. coli* adherence factors as the bacteria adapted to an intracellular lifestyle (2) due to three main reasons. First, *Shigella* grown in standard laboratory media lack structures upon transmission electron microscopy (TEM) (17, 18), unlike some strains of *E. coli* in which adherence factors are thought to be constitutively expressed (19, 20). Second, examination of *Shigella* genomes deposited in GenBank reveals that almost all adherence gene clusters, such as type 1 fimbriae (10, 21) and curli (22), contain at least one annotated pseudogene that is crucial for either the adherence factor structure or the assembly process (17, 23, 24). Third, production of adherence factors is considered counterproductive to the lifestyle of an intracellular pathogen evading immune detection (2, 25, 26).

Despite this null adherence factor hypothesis, a few reports have detected adherence factor expression in *S. flexneri* (27–29); but in-depth genetic analyses were not performed. Furthermore, we have previously demonstrated that tryptic soy broth media supplemented with bile salts induce the adherence of *S. flexneri* 2457T to colonic epithelial cells, which is facilitated at least in part by the type-III secretion system effector proteins OspE1 and OspE2 (30). Finally, our recent publication characterizes an adhesive biofilm phenotype following prolonged exposure to a combination of bile salts and glucose that represent aspects of the *in vivo*-like conditions (IVLCs) found in the small intestine (31). Given this literature and the fact that deletion of both *ospE1* and *ospE2* did not completely abrogate adherence (30), we sought to determine if additional adherence factors were produced by *S. flexneri* 2457T following IVLC exposure. In the first part of our analysis, we utilized electron microscopy (EM) to confirm the presence of putative adherence factors following IVLC exposure, and subsequently characterized the transcription profile of the annotated adherence gene clusters. In the second part of our analysis, we performed mutational and proteomic analyses to characterize three of the adherence structural genes and demonstrated that these factors facilitate adherence for both biofilm formation and colonization of colonic epithelial cells, particularly in the human intestinal organoid-derived epithelial monolayer (HIODEM) model. This work broadens our understanding of *S. flexneri* 2457T pathogenesis and demonstrates that *S. flexneri* 2457T likely expresses a number of “traditional” adherence factors important for pathogenesis. Insights gained from this work could have an important impact on *Shigella* therapeutic and vaccine development.

## Results

### *S. flexneri* 2457T produces putative adherence structures in IVLCs

Previous studies have demonstrated that *S. flexneri* 2457T grown in IVLCs produced a biofilm. Furthermore, upon bacterial dispersion from the biofilm, recovered bacteria displayed induced adherence to colonic HT-29 cells. This analysis enabled us to expand the *Shigella* infection paradigm to incorporate biofilm formation during small intestinal passage, dispersion upon colonic transition following the loss of the bile salts signal, and induced infection rates due to the IVLCs encountered in the small intestine (31, 32). Since adherence factors are important components of biofilm formation (32, 33), we performed EM analysis of bacteria isolated from the IVLC-induced biofilm to visualize possible adherence factors. As shown in Figure 1, bacteria produced thick appendages; thinner, hair-like appendages; and electron dense, cloud-like aggregates in IVLCs. Bacteria grown in Luria broth (LB) and LB supplemented with glucose (2% w/v) lacked structures while bacteria grown in LB supplemented with bile salts (0.4%) produced very minimal structures. Utilization of bile salts in tryptic soy broth (TSB) media, in which there is added glucose relative to LB (31), resulted in a similar appearance of putative adherence factors as in LB media supplemented with both glucose and bile salts (**Supplemental Figure S1**). The data confirmed our observations that glucose and bile salts (IVLCs) are required for *S. flexneri* 2457T to form an adhesive biofilm (31). To support the biofilm data and our previous induced HT-29 adherence observations (30, 31), we performed adherence analysis on a human intestinal organoid-derived epithelial monolayer (HIODEM) model. The model is derived from stem cells isolated from intestinal tissue, propagated as organoids, and subsequently trypsinized and seeded onto transwells to generate a two-dimensional (2-D) polarized, differentiated model of the intestinal epithelium in which enterocytes, mucus-producing goblet cells, and antigen sampling M cells are present (34–38). With the model derived from the ascending colon, *S. flexneri* 2457T subcultured in IVLCs displayed putative adherence factors contacting the epithelial cells (Figure 2). In all, the data suggested these putative adherence factors were important for both biofilm formation and adherence to colonic epithelial cells.

**Figure 1.**
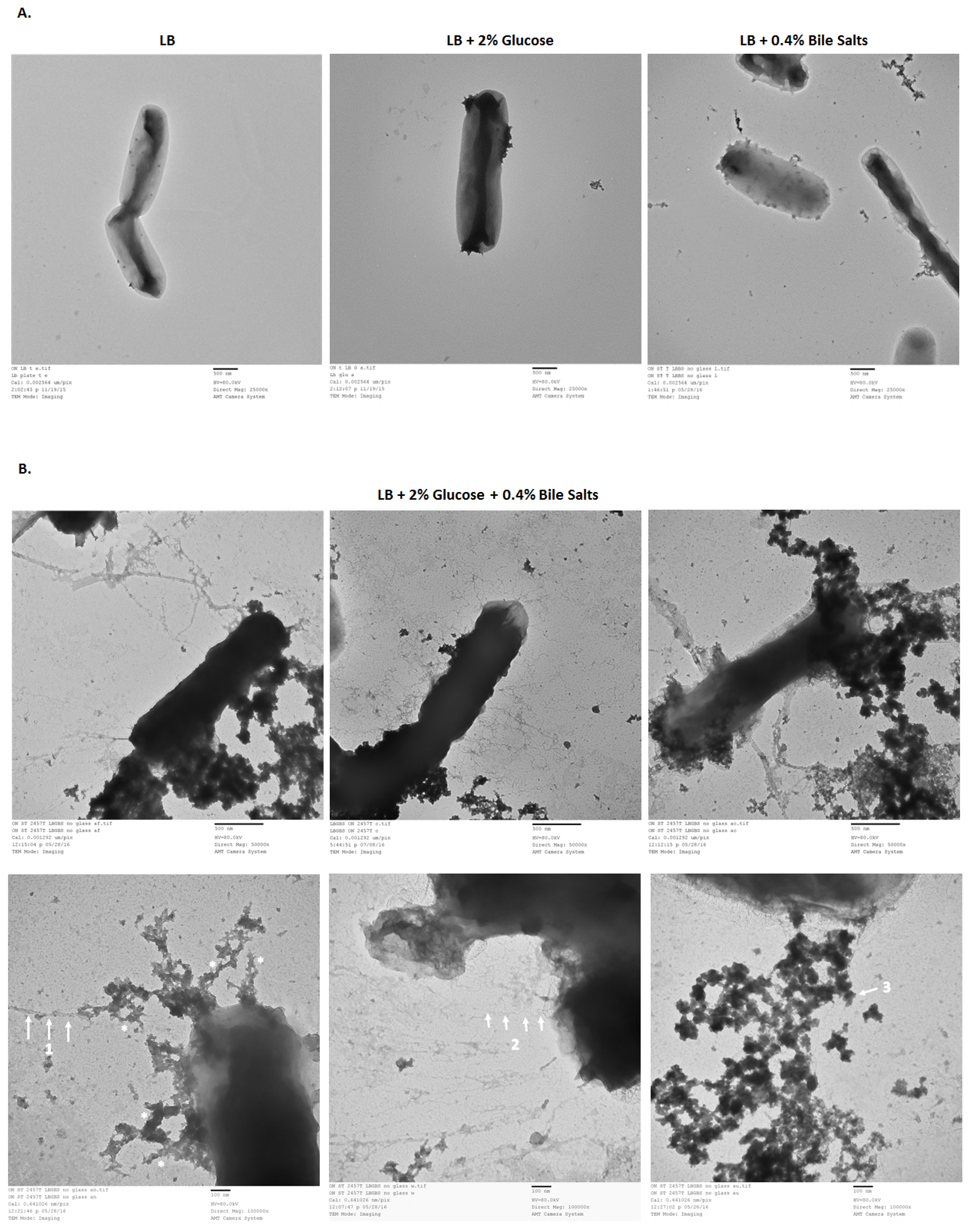
Growth of *S. flexneri* 2457T in IVLCs produces several putative adherence factors. Wild type cultures of 2457T were grown overnight with static growth in the indicated media. Samples were negatively stained and imaged by electron microscopy. Images are representative of at least 3 biological replicates. **A.** TEM analysis of 2457T grown in LB, LB supplemented with 2% glucose, or LB supplemented with 0.4% bile salts demonstrated that adherence factors were either not produced or minimally produced in these conditions (all images are at 25,000 X magnification, scale bar 500 nm). **B.** 2457T grown in LB supplemented with both 2% glucose and 0.4% bile salts (IVLCs) revealed three types of putative adherence factors upon TEM analysis. Top row of images is 50,000 X magnification (500 nm scale bar) and bottom row of images is 100,000 X magnification (100 nm scale bar). Arrow 1 points to thicker appendages; arrow 2 points to thinner, hair-like appendages; and arrow 3 points to electron dense, cloud-like aggregates. The asterisks (*) denote rough, complex structures (please refer to discussion).

**Figure 2.**
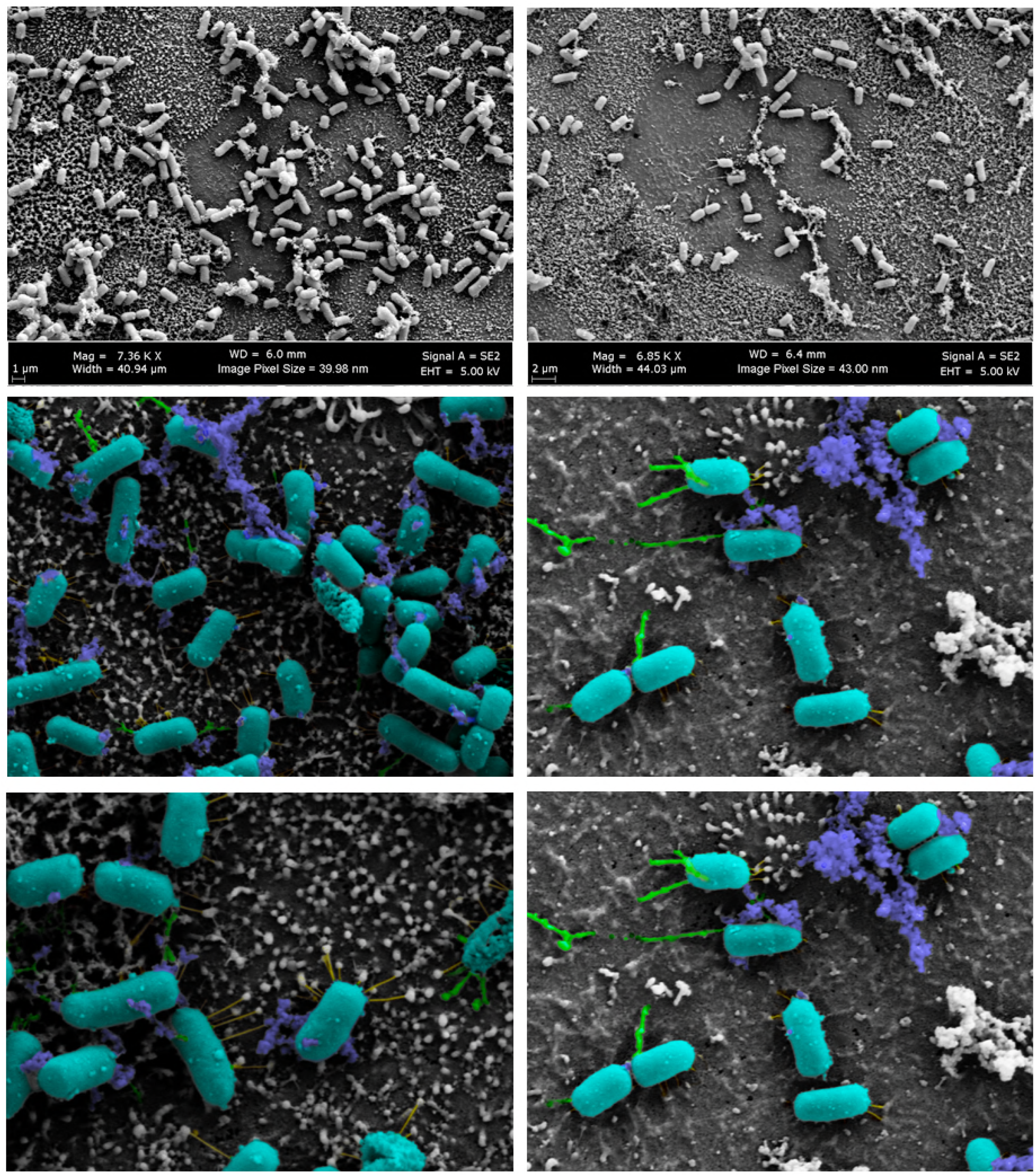
Scanning electron microscopy of *S. flexneri* 2457T adherence on the human intestinal organoid-derived epithelial monolayer model. *S. flexneri* 2457T was subcultured in IVLCs (TSB + bile salts), washed, and applied for adherence analysis on the HIODEM model. Following infection, samples were fixed and processed for SEM analysis. The three types of putative adherence factors are visualized on the bacteria interacting with the apical surface of the epithelial cells. Pseudo-coloring was performed to enhance visualization of the different structures. The bacterial rods were colored teal, thick appendages were colored green, thin hair-like projections were colored yellow, and cloud-like aggregates were colored blue. Left and right columns represent two separate biological samples. Magnification ranges from approximately 7,000 X (top) to approximately 20,000 X (center and bottom).

### *S. flexneri* 2457T maintains and transcribes several adherence gene clusters

We next examined the transcription of the *S. flexneri* 2457T adherence genes under various conditions. *In silico* analyses of the annotated *S. flexneri* 2457T genome on NCBI GenBank identified several adherence gene components (Table 1, Figure 3, and **Supplemental Figure S2**). These genes are maintained in *S. flexneri* 2457T despite examples of full gene and/or operon deletions for some of the adherence gene clusters in other *Shigella* species (23). As documented in previous studies (17, 23, 24), all *S. flexneri* 2457T adherence gene clusters contain at least one annotated pseudogene (due to predicted point mutations, truncations, or insertion sequences) that support hypotheses stating that *Shigella* cannot produce “traditional” adherence factors. However, previous RNA-seq data (31) indicated that despite the gene annotations, most of the adherence genes were transcribed by *S. flexneri* 2457T (Figure 3 and **Supplemental Figure S2**). To confirm the RNA-seq results, we performed RT-PCR analysis of the annotated adherence gene clusters (Figure 3 and **Supplemental Figure S2**). RNA isolated from *S. flexneri* 2457T broth cultures were positive for transcription of the adherence genes and large segments of the predicted operons. Insertion sequences did not prevent transcription of large downstream segments. For example, as demonstrated in Figure 3, we amplified cDNA products from *fimD* just after the insertion sequences to the end of *fimF*. Finally, we utilized quantitative RT-PCR analysis for additional data to support the transcription of adherence genes. As described in previous literature for other pathogens (39–42), glucose induced the expression of the *S. flexneri* 2457T genes encoding structural subunits (Figure 4). In all, the data indicate that adherence gene clusters are genomically maintained and transcriptionally regulated in *S. flexneri* 2457T despite the pseudogene annotations.

**Figure 3.**
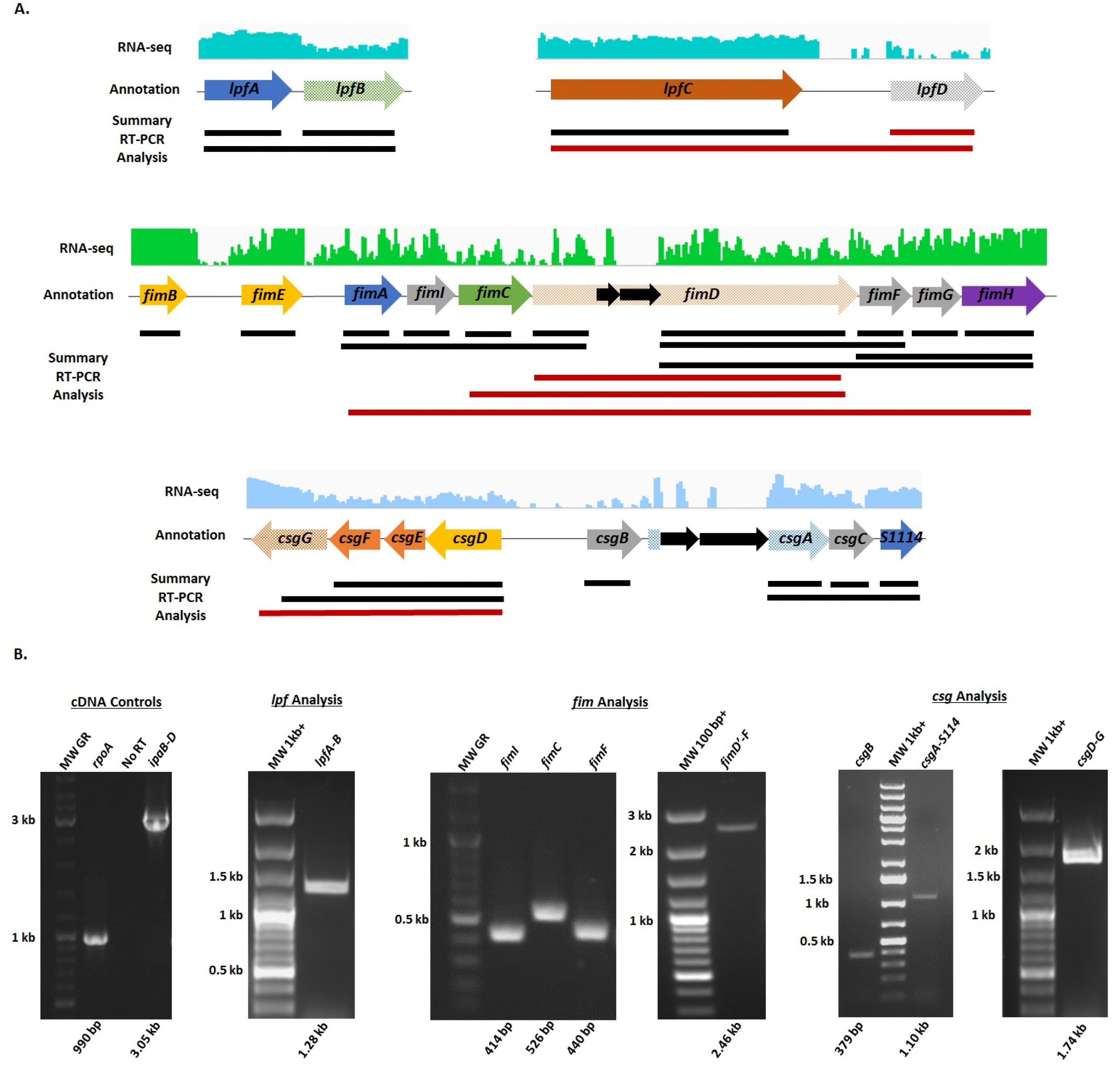
*In silico* and transcriptomic analysis of the *S. flexneri* 2457T adherence gene clusters. **A.** Identification of the long polar fimbriae *(lpf)*, type 1 fimbriae (fm), and curli *(csg)* adherence gene clusters in the *S. flexneri* 2457T genome was performed using the NCBI GenBank tool. Arrows represent annotated open reading frames (ORF) in which blue arrows represent the annotated major subunits, green arrows represent the annotated chaperones, orange arrows represent the annotated ushers or pores, yellow arrows represent the annotated regulatory subunits, purple arrows represent the annotated tip adhesins, and grey arrows represent additional putative adherence genes involved in assembly or secretion. All arrows with checkered backgrounds are annotated as pseudogenes in NCBI due predicted insertion sequences, truncations, or frameshifts. Insertion sequences or transposons are represented by solid black arrows. The RNA-sequencing trace read data are provided for each gene cluster above the arrows. The best trace read available is presented, with green representing shaking growth conditions (darker green in bile salts) and blue representing static growth conditions (darker blue in bile salts). Solid lines below the arrows represent the summary of confirmation of gene transcription that resulted from non-quantitative RT-PCR analyses. Single gene and polycistronic amplifications were performed to obtain the largest products possible for each operon. Red lines represent products that were not obtained. Please refer to Supplemental Figure 2 for additional gene cluster analyses. **B.** Representative gel electrophoresis images of non-quantitative RT-PCR analyses of the various adherence gene clusters are provided. Biologically independent RNA samples were used in the analysis in which RNA integrity was verified by amplifying the house-keeping gene *rpoA* as well as the *ipaB* to *ipaD* operon encoded on the virulence plasmid. Control amplifications without reverse transcriptase in the *rpoA* reactions ensured there was no DNA contamination of the RNA samples prior to cDNA synthesis. Each gene is labeled with the expected molecular weight (MW) of the product provided below the gel image. Note that different MW ladders were utilized in the analyses. Please refer to the Materials and Methods section for more information.

**Figure 4.**
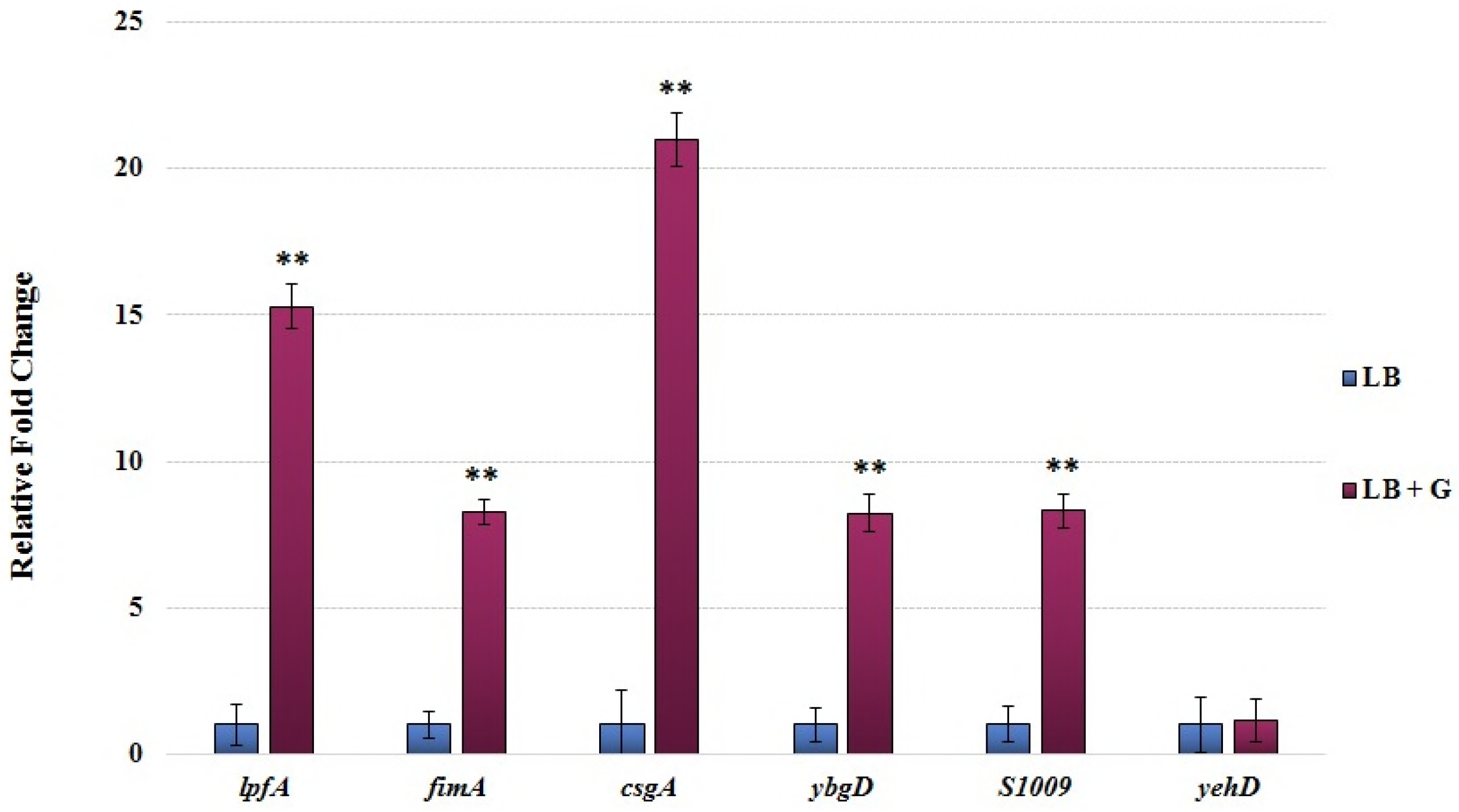
Quantitative RT-PCR analysis of *S. flexneri* 2457T structural adherence genes. RNA was isolated from *S. flexneri* 2457T grown in media (LB) or media supplemented with 2% w/v glucose (LB + G). For each primer set the relative fold change +/− the standard error of the ΔCt value is plotted for each gene. Data represent the average of at least three biological independent samples in which each sample had technical duplicates. *yehD* (see Table 1) expression was not induced in the presence of glucose; and therefore, served as an internal negative control for the analysis. ** denotes significance in which all p-values are ≤ 0.01 for each gene between LB and LB + G conditions.

**Table 1.** Adherence gene clusters identified in the *Shigella flexneri* 2457T genome.

| Adherence Gene | Locus Tag | Annotated Function/Pseudogene |
| --- | --- | --- |
| <u>Long Polar Fimbriae</u> |  |  |
| *lpfA* | **S3961** | **Major fimbrial subunit, deleted in this study** |
| *lpfB* | - | Chaperone/Pseudogene |
| *lpfC* | S4048 | Outer membrane usher |
| *lpfD* | - | Fimbrial protein/Pseudogene |
| <u>Type 1 Fimbriae</u> |  |  |
| *fimB* | S4467 | Recombinase |
| *fimE* | S4466 | Tyrosine recombinase |
| *fimA* | **S4465** | **Major fimbrial subunit, deleted in this study** |
| *fimI* | S4464 | Pilus biosynthesis protein |
| *fimC* | S4463 | Chaperone |
| *fimD* | S4462 | Outer membrane usher/Pseudogene |
| *fimF* | S4458 | Minor subunit |
| *fimG* | S4457 | Minor subunit |
| *fimH* | S4456 | Tip adhesin |
| <u>Curli Operon</u> |  |  |
| *csgG* | S1104 | Assembly protein/Pseudogene |
| *csgF* | S1105 | Assembly protein |
| *csgE* | S1106 | Assembly protein |
| *csgD* | S1107 | Transcriptional regulator |
| *csgB* | **S1108** | **Minor subunit, deleted in this study** |
| *csgA* | **S1109** | **Major subunit/Pseudogene, deleted in this study** |
| *csgC* | S1113 | Autoagglutination assembly protein |
| S1114 | S1114 | Hypothetical (*fimA* homolog) |
| <u>ybg Operon</u> |  |  |
| *ybgD* | S0591 | Fimbrial-like protein (*fimA* homolog) |
| *ybgQ* | S0592 | Outer membrane protein (*fimD* homolog) |
| *ybgP* | S0593 | Chaperone (*fimC* homolog) |
| *ybgO* | S0594 | Putative fimbrial-protein/Pseudogene |
| <u>ycb Operon</u> |  |  |
| *ycbQ* | S1003 | Fimbrial-like adhesin/Pseudogene |
| *ycbS* | S1006 | Outer membrane usher protein |
| S1007 | S1007 | Fimbrial protein |
| S1008 | S1008 | Fimbrial-like protein |
| S1009 | S1009 | Fimbrial-like protein (*fimA* homolog) |
| *ycbF* | S1010 | Putative pili assembly chaperone |
| <u>yeh Operon</u> |  |  |
| *yehD* | S2298 | Fimbrial-like protein |
| *yehC* | S2297 | Chaperone/Pseudogene |
| *yehB* | S2296 | Outer membrane usher protein |
| *yehA* | S2295 | Putative fimbrial-protein/Pseudogene |
| <u>yra Operon</u> |  |  |
| *yraH* | S3396 | Fimbrial-like adhesin/Pseudogene |
| *yraI* | S3397 | Chaperone (*fimC* homolog) |
| *yraJ* | S3398 | Outer membrane usher/Pseudogene |
| *yraK* | S3403 | Fimbrial Protein |
| <u>S4250-S4254</u> |  |  |
| S4250 | S4250 | Pili assembly chaperone (*papD_C* homolog) |
| S4254 | S4254 | Outer membrane usher/Pseudogene |
| <u>S3342-S3341</u> |  |  |
| S3342 | S3342 | Hypothetical (*fimD* homolog) |
| S3341 | S3341 | Hypothetical (pilus biogenesis initiator homolog) |
| <u>sfm Operon</u> |  |  |
| *sfmA* | S0469 | Fimbrial-like protein (*fimA* homolog) |
| *sfmC* | S0470 | Chaperone |
| *sfmD* | - | Outer membrane usher/Pseudogene |

### Mutational analyses of adherence structural genes

We next performed mutational analyses of the genes encoding major structural subunits to demonstrate functional roles in biofilm formation and epithelial cell adherence. We concentrated our analyses on long polar fimbriae representing the thick appendages, type 1 fimbriae representing the thin hair-like structures, and curli representing the electron dense, cloud-like aggregates based on our *in silico* analyses, the combined appearance of structures in Figures 1 and 2, and the known functional roles of these structures in initial biofilm formation and epithelial cell adherence in other pathogens (8, 28, 43–45). Thus, we constructed *ΔlpfA, ΔfimA*, and *ΔcsgA* mutants. For curli, we also constructed a double *ΔcsgAB* mutant due to the additional role of the *csgB* minor subunit in adherence (46). The mutants were subsequently evaluated by EM for loss of recognized surface structures. Each mutation resulted in loss of the predictive adherence factor structure while also facilitating visualization of the two other predominant structures (Figure 5A). Furthermore, ammonium sulfate precipitation for the isolation of adherence factors (47) was performed to verify our results and enabled visualization of adherence factors in both wild type and mutant strains (Figure 5B). Finally, to confirm the presence of curli despite the disorganized appearance, the Congo red binding assay was performed given the ability of Congo red dye to bind the amyloid structures of curli and produce a birefringence signal under polarized light (48–50). A positive birefringence signal was detected for both wild type 2457T and the *ΔlpfA* mutant, which was used as a mutation control for this assay. However, the *ΔcsgA* mutant did not have a birefringence signal, indicating that the mutation resulted in loss of curli production (**Supplemental Figure S3**). In all, these analyses suggested we identified genes encoding the structural subunits of the putative adherence factors expressed by *S. flexneri* 2457T.

**Figure 5.**
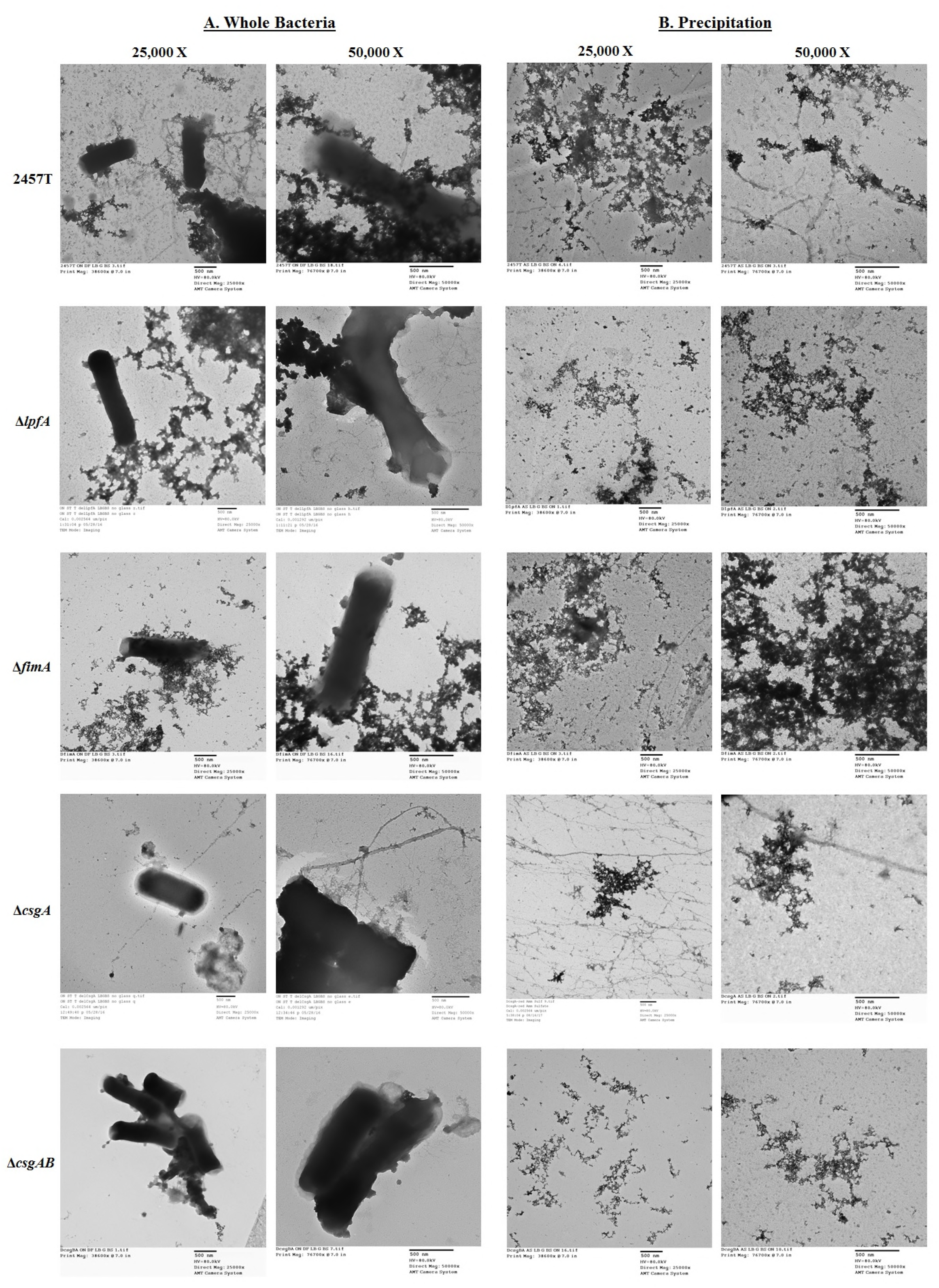
TEM analysis of the *S. flexneri* 2457T mutants. **A.** The *ΔlpfA, ΔfimA*, ΔcsgA, and *ΔcsgAB* were grown statically overnight in IVLCs (LB+ 2% glucose + 0.4% w/v bile salts), subsequently processed for TEM analysis, and analyzed with wild type 2457T as a control. Each mutation resulted in the loss of either the thick appendages *(ΔlpfA)*, the thinner, hairlike appendages *(ΔfimA)*, or the electron dense, cloud-like aggregates (ΔcsgA, and ΔcsgAB). Images for wild type 2457T are from separate, biologically independent experiments relative to the images provided for Figure 1. **B.** To verify the results, ammonium sulfate precipitation was performed to isolate and visualize the adherence factors from wild type *S. flexneri* 2457T and each of the four mutants. The three types of factors can be visualized in wild type bacteria; however, only two of the three structures were present for the mutants. Each mutation resulted in the expected loss of structure. The data verify the correct structural subunit was deleted for each mutant. All images are representative of at least two biological independent experiments. Different fields are presented for the 25,000X and 50,000X magnifications for all images in (A) and (B).

### Functional analyses of the adherence mutants

Functional analyses of the mutants were next performed to evaluate the role of each factor in biofilm formation and adhesion to epithelial cells. First, given the importance of adherence in the initiation of biofilms (33, 51), we analyzed the mutants in the biofilm assay (31, 32). All mutants exhibited reduced biofilm formation at 3 hours (Figure 6), a timepoint used to examine the role of adherence factors in early biofilm formation (32). Thus, we concluded that structures encoded by *lpfA, fimA*, and *csgAB* have roles in the adhesion process for IVLC-induced biofilm formation in *S. flexneri* 2457T.

**Figure 6.**
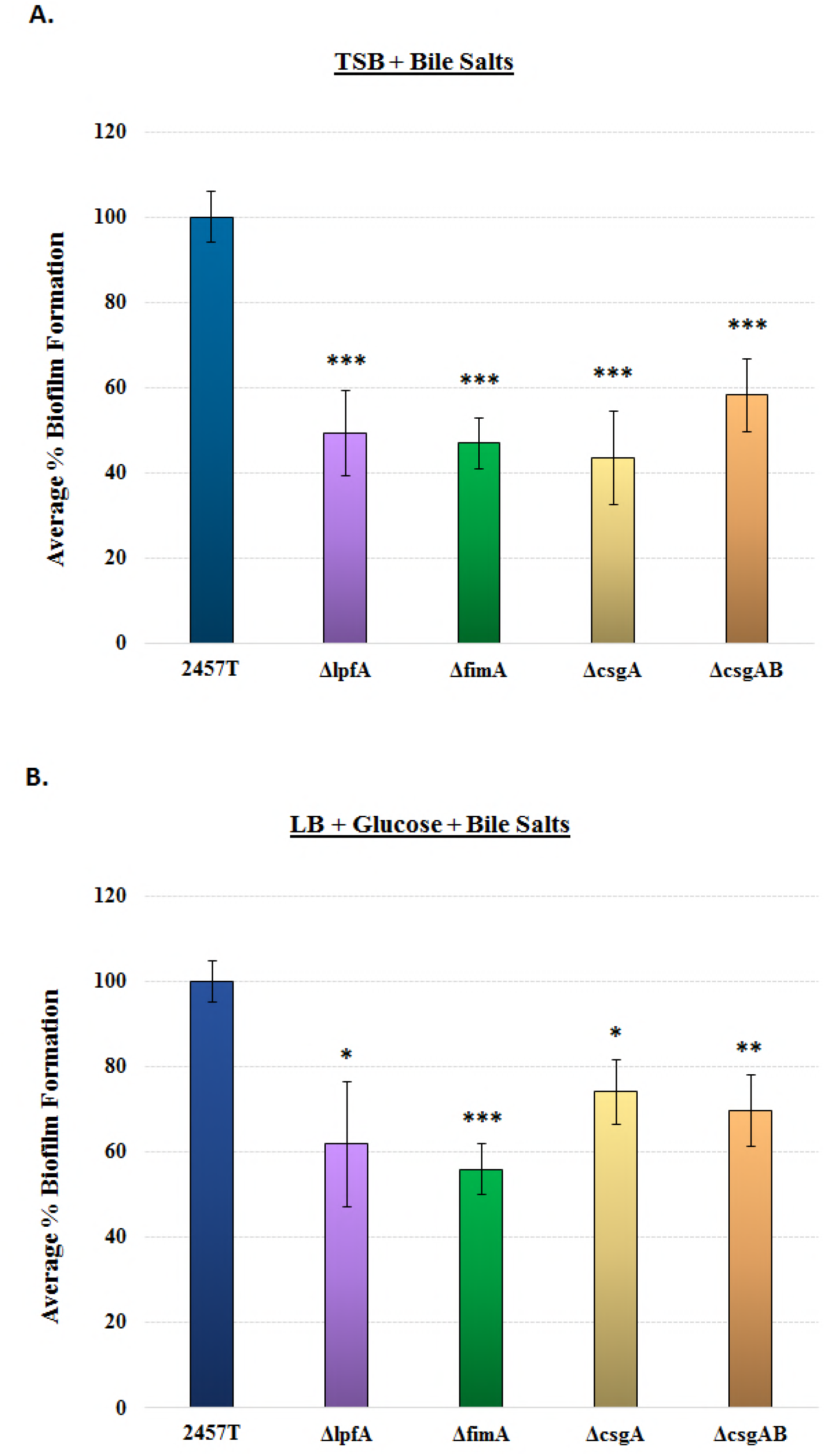
Analysis of biofilm formation for each *S. flexneri* 2457T adherence mutant. Wild type *S. flexneri* 2457T and the adherence mutants *(ΔlpfA, ΔfimA*, ΔcsgA, and *ΔcsgAB)* were analyzed for biofilm formation at 3 hours post-inoculation to examine the adherence phase of biofilm formation. Each mutant had a significant reduction in biofilm formation. The average percent (%) biofilm formation +/− standard error relative to 2457T is plotted for each mutant. All data represent three biological independent experiments in which each experiment had technical triplicates. The p-values ranged from <0.05 (*) to <0.01 (**) and <0.001 (***). **A.** Assays performed in TSB media supplemented with bile salts. **B.** Assays performed in LB media supplemented with glucose and bile salts.

Second, we analyzed adherence to epithelial cells. We hypothesized that the adherence factors expressed in the IVLCs would also facilitate epithelial cell contact. This hypothesis was supported by our previous observations of induced *S. flexneri* 2457T adherence to HT-29 cells following biofilm dispersion in conditions that mimicked the loss of the bile salts signal during the terminal ileum to colon transition (31). As with the biofilm assay, all mutants had significant reductions in adherence relative to wild type bacteria, with the Δ*fimA* and *ΔcsgAB* mutants having the greatest reductions (Figure 7A). The double *ΔospE1/ospE2* mutant (BS808) served as an adherence mutant control given our previous analysis of the role of OspE1 and OspE2 in bile salt-mediated adherence (30). To ensure the mutations did not affect the overall invasive ability of each strain, invasion assays were performed using conventional methods of centrifugation to initiate host cell contact (52). All mutants retained wild type levels of invasion following centrifugation of the bacteria onto the HT-29 cells (data not shown), which confirmed that the mutations did not affect the basic invasion phenotype of the strains. Finally, to confirm the HT-29 adherence data, we evaluated the *ΔlpfA, ΔfimA*, and *ΔcsgA* mutants in the HIODEM model and found that each mutant had significantly reduced adherence relative to wild type bacteria (Figure 7B). EM analysis of infected samples enabled visualization of mutants with a smoother surface and less adherence factors relative to wild type bacteria (Figure 7C, Figure 2). In all, the data demonstrated that factors encoded by *lpfA, fimA*, and *csgAB* in *S. flexneri* 2457T have a functional role in adherence to colonic epithelial cells.

**Figure 7.**
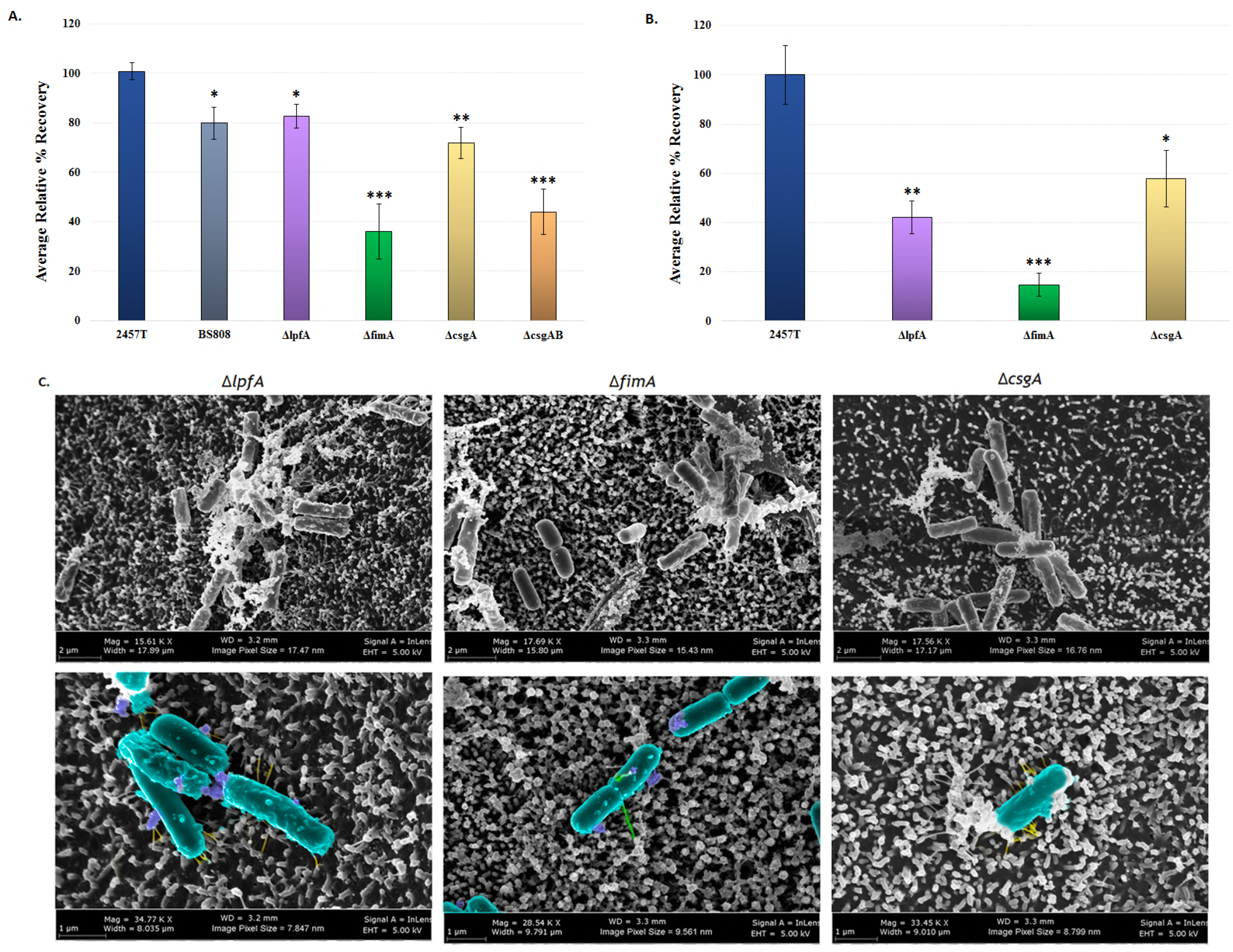
Analysis of adherence with the mutant strains. **A.** Wild type *S. flexneri* 2457T, the control BS808 *(ΔospE1* + ΔospE2), and the adherence mutants *(ΔlpfA, ΔfimA*, ΔcsgA, and *ΔcsgAB)* were grown overnight as described for the biofilm assay. On the next day, bacteria were collected, washed with 1X PBS, and analyzed for adherence to HT-29 cells. Each mutant had a significant reduction in adherence relative to wild type 2457T, with the most significant reductions seen with the *ΔfimA* and *ΔcsgAB* mutants. **B.** To verify the HT-29 data, the *ΔlpfA, ΔfimA*, and *ΔcsgA* mutants were evaluated for adherence in the HIODEM model. Like with the HT-29 cells, each mutant displayed a significant reduction in adherence relative to wild type 2457T, with the *ΔfimA* mutant having the greatest reduction. For both A and B, the average percent (%) recovery of adherent bacteria +/− standard error relative to wild type 2457T is plotted. The HT-29 infections represent three independent experiments in which each experiment had technical triplicates. The HIODEM infections represent two independent experiments in which each experiment had at least two technical replicates. The *p*-values ranged from <0.05 (*) to <0.01 (**) and <0.001 (***). **C.** Scanning electron microscopy analysis of the *ΔlpfA, ΔfimA*, and *ΔcsgA* mutants on the HIODEM model enabled visualization of the reduced adherence factors on each mutant. The *ΔlpfA* mutant lacks the thick appendages, the *ΔfimA* mutant lacks the thin hair-like projections, and the *ΔcsgA* mutant lacks the cloud-like aggregates. Magnification ranges from 15,000 X to 35,000 X. Pseudo-coloring was performed as described for Figure 2.

### Mass spectrometry analysis to evaluate secretion of the adherence structural proteins in IVLCs

As a final method to confirm the presence and functional secretion of LpfA, FimA, and CsgAB structural proteins, proteomic analyses were performed on culture supernatants from the biofilm assay. Both intact mass spectrometry (MS) analysis and peptide fingerprinting MS/MS analysis of trypsin-digested samples confirmed the presence of LpfA, FimA, CsgA, and CsgB, with each protein having high levels of sequence coverage upon the fingerprinting MS/MS analysis (Table 2), verifying that the proteins are secreted in IVLCs. Due to the complexity of the samples for MS analysis, especially from the extracellular polymeric substance (EPS) matrix production of the IVLC-induced biofilm (31), a higher than expected mass error was observed. Therefore, we cloned the *lpfA, fimA*, and *csgA* genes from *S. flexneri* 2457T, added a histidine tag to the genes, and transformed each respective mutant to perform immunoprecipitation and complementation analyses. As shown in Figure 8, the tagged LpfA, FimA, and CsgA proteins were expressed in the respective mutants, secreted, and purified from IVLC-induced biofilm culture supernatants, which verified the MS data. Biofilm assay analyses and TEM visualization of the over-expressed structures verified these tagged constructs were functional (Figure 8). In all, the data confirmed the EM and mutation analyses, and verified that *lpfA, fimA*, and *csgAB* genes produce functional proteins in *S. flexneri* 2457T.

**Figure 8.**
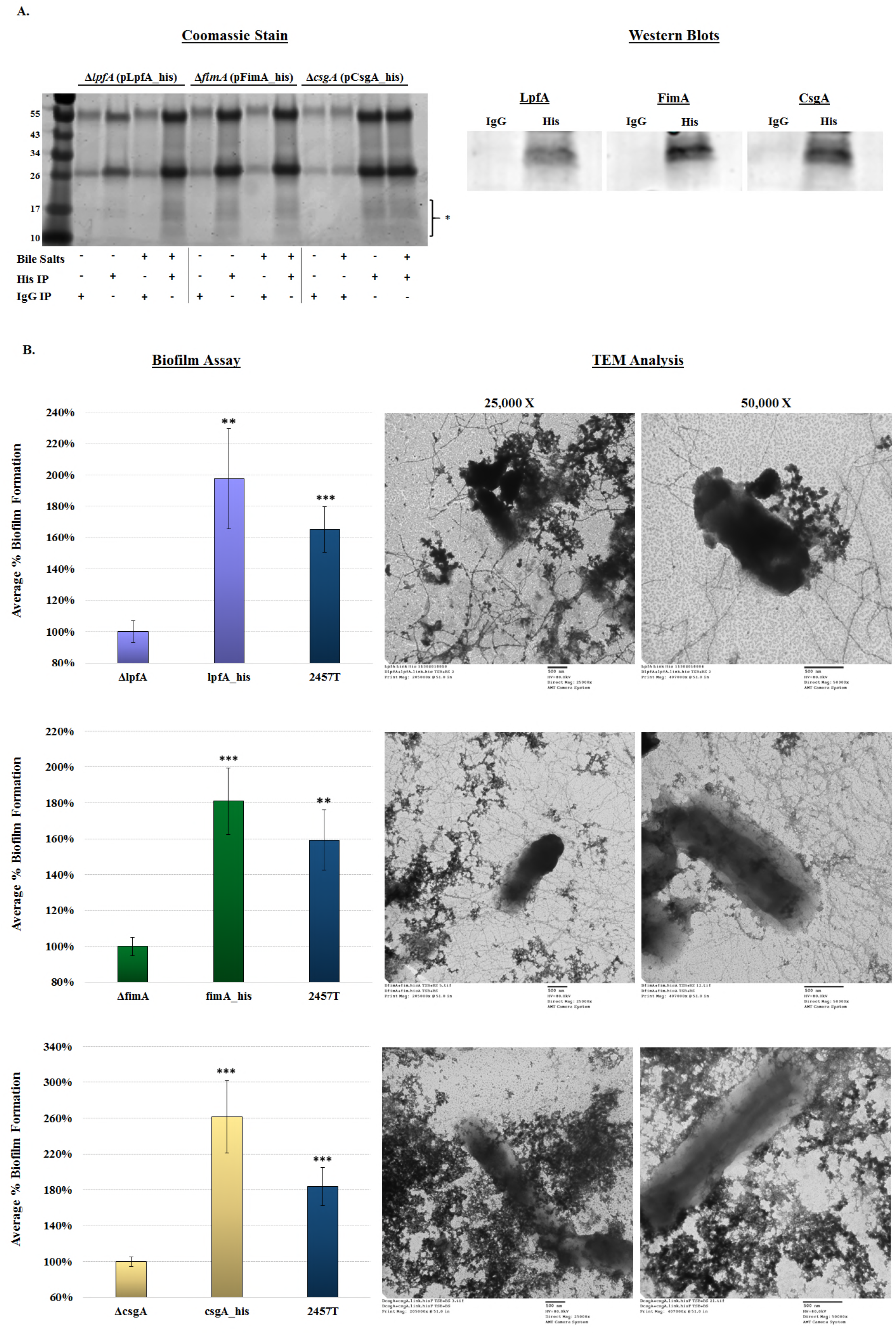
Immunoprecipitation and functional complementation of the histidine-tagged proteins. **A.** To confirm the mass spectrometry analysis, bacteria expressing his-tagged proteins were cultured in the overnight biofilm assay and the culture supernatants were immunoprecipitated. Left: IgG negative control and anti-his immunoprecipitation experiments in TSB +/− bile salts were analyzed by SDS-PAGE with Coomassie blue staining. Proteins in the 10 to 17 kD range (noted by bracket and *) were immunoprecipitated with the anti-his resin and in the presence of bile salts. Right: Western blot analysis of proteins in the 10-17 kD range confirmed the his-tagged LpfA, FimA, and CsgA were secreted into the supernatant of the biofilm culture. No proteins were detected in the negative control samples in which IgG was used in the immunoprecipitation. **B.** Biofilm and TEM analyses were used to verify that the his-tagged proteins were functional and could complement the respective mutants. The complemented strains (*lpfA+* (top), *fimA+* (middle), and *csgA+* (bottom)) produced significantly higher biofilms relative to the mutants to indicate restoration of function, while TEM analysis enabled visualization of the over-expressed structures. The biofilm assays were analyzed at 4.5 hours. The data represent the average of two independent experiments, each with six technical replicates, +/− standard error. All p-values ranged from <0.01 (**) to <0.001 (***). Each complement averaged the same or above wild type 2457T levels (differences not significant).

**Table 2.** Mass spectrometry analysis of the bile salt-induced biofilm supernatants.

| Protein | Intact Average<br>Mass Observed<br>(Da) | Intact Average<br>Mass Expected<br>(Da) | Mass Error<br>(PPM) | Response<br>(Total Ion Count) | Digested Peptide<br>Coverage |
| --- | --- | --- | --- | --- | --- |
| LpfA | 19,644.38 | 19,642.80 | 86.8 | 583,652,552 | 100% |
| FimA | 18,348.20 | 18,347.28 | 49.8 | 221,292,058 | 82% |
| CsgA | 12,389.58 | 12,388.92 | 53.4 | 203,325,206 | 84% |
| CsgB | 15,870.81 | 15,869.62 | 74.9 | 369,128,136 | 91% |

## Discussion

Characterization of the three structural genes in this study demonstrates that *S. flexneri* 2457T utilizes “traditional” adherence factors to initiate biofilm formation and to facilitate contact to colonic epithelial cells. Several observations influenced the investigation, including the lack of an adherence null mutant in OspE1 and OspE2 analysis (30), the subsequent biofilm formation and induced adherence observed following IVLC exposure (31, 32), as well as the presence of the various adherence gene clusters in the *S. flexneri* 2457T genome. The literature on “traditional” *Shigella* adherence factors is contradictory. Numerous studies have suggested that the various gene clusters have been lost during evolution as a pathoadaptive response to the host. Notably, laboratory growth methods consistently used to demonstrate fimbrial production in strains of *E. coli* (19, 20) were not successful for either lab strains and clinical isolates of *Shigella* (17, 18). Our control media analyses, in which the combination of glucose and bile salts were absent, confirmed many of these previous findings on the phenotypic level. The visualization of putative adherence factors required the addition of both glucose and bile salts to the media, factors that are present in the small intestine during host transit (31, 32, 53, 54). Interestingly, glucose induces the transcription of the structural subunits (Figure 4), yet adherence factors were not visible in LB + glucose treatment while minimal adherence factors were visualized in the LB + bile salts treatment (Figure 1). Thus, based on the data presented in Figures 1 and 4, we hypothesize that glucose induces structural gene transcription while bile salts serve as a secretion signal. The amount of glucose required for signaling can vary, as evident by the different percentages of glucose in TSB compared the glucose-supplemented LB, which is consistent with our previous observations (31). Nevertheless, this work demonstrates that *S. flexneri* produces these factors, while also highlighting the importance of using physiologically relevant conditions to study bacterial pathogenesis, especially for human-adapted pathogens like *Shigella*.

The combined RNA-seq and RT-PCR analyses of the adherence gene clusters demonstrate that some of the gene annotations are accurate, while other annotations require refinement. For example, the *csgG* gene is annotated as a pseudogene due to a point mutation that creates an in-frame stop codon. The RT-PCR analysis confirmed this annotation since a partial *csgG* product was detected prior to the stop codon; however, no product was detected with a reverse primer that annealed downstream of this mutation. As another example, there was significant transcription of the *ycbQ* gene despite the truncated pseudogene annotation. Finally, while the full *ybgO* gene could not be amplified under the conditions examined, inspection of the primary genomic sequence (GenBank Accession Number S0594) combined with the RNA-seq read mapping indicate that two separate open reading frames or small RNAs may be transcribed in this region. The effects of transcription of these partial gene fragments on *S. flexneri* 2457T gene regulation or adherence factor expression will require additional analyses.

The mutational and complementation analyses demonstrate functional roles for long polar fimbriae encoded by *lpfA* structural gene, type 1 fimbriae encoded by *fimA* structural gene, and curli encoded by the *csgA* and *csgB* structural genes. Long polar fimbriae have been shown to be important for pathogenic *E. coli* and *Salmonella* interactions with M cells during intestinal colonization, and the *lpfA* genes have been demonstrated to be induced by bile salts (13, 55–58). As seen in Figure 2, thicker appendages are bound to the surface of cells lacking microvilli, which is a hallmark of M cells (59). Additionally, the *ΔlpfA* mutation had a greater effect on adherence in the HIODEM model in which M cells are present (34–37) compared to HT-29 cells alone (Figure 7). For type 1 fimbriae, previous studies support our observations of both *fimA* gene transcription and soluble FimA expression. First, clinical isolates of *Shigella* produced fimbrial-like adhesins after periods of prolonged static growth; however, the genes encoding the factors were not identified (28). Second, another RNA-seq study detected significant induction of the type 1 *fim* operon in a *ΔicgR* mutant of *S. flexneri* 2457T during the intracellar phase of the *Shigella* lifestyle (60). Finally, soluble *S. flexneri* FimA protects mitochondrial integrity and epithelial cell survival during infection (61). It is worth noting that the predicted type 1 fimbrial-like structures visualized from the biofilm assays (Figures 1 and 5) appear thinner compared to the fimbrial-like structures visualized during infection (Figures 2 and 7). We do not expect the structures to appear like typical observed *E. coli* structures, especially since a truncated or substituted FimD (see below) could affect assembly. While the *ΔfimA* mutant analyses resulted in less visualized fimbrial-like structures (Figures 5 and 7), we currently cannot rule out the contribution of or compensation by the additional *S. flexneri* 2457T *fimA* homologs (Table 1), particularly in bile salt conditions that induce such a strong biofilm response (31, 62). As with *lpfA* and *csgA*, TEM analysis of the histidine-tagged *fimA^+^* complement verified the appearance of the structures (Figure 8). Thus, we have provided strong evidence that the type 1-like fimbriae visualized in our analyses is due to expression from the *fimA* structural gene.

The curli in *S. flexneri* 2457T appears disorganized compared to the conventional fiber structures detected in other pathogens (43, 48). This lack of assembly could be due to the fact that CsgA is truncated or due to the incomplete production of CsgG, the outer membrane lipoprotein involved in the stability of the curlin proteins during assembly (22, 43, 63). Furthermore, a truncated CsgG may prevent appropriate interaction with CsgF, thereby affecting curli assembly (43, 64). Our analyses indicate a soluble portion of CsgA is produced in *S. flexneri* 2457T that is sufficient to provide function in adherence, particularly in the establishment of the IVLC-induced biofilm. This soluble portion of the CsgA protein is likely facilitated by a functional CsgB minor subunit protein given the further reduction in phenotypes of the double *ΔcsgAB* mutant, the visualization of electron dense aggregates in the *ΔcsgA* mutant (Figures 5 and 7), and the demonstration that CsgB has a role in adherence (46). Interestingly, our EM images suggest that the curli subunits exploit other adherence structures as a scaffold for more appropriate organization *(e.g*., see the rough, complex structures marked by asterisks in Figure 1B). Moreover, the additional electron dense material visualized in the curli mutants, particularly with Δ*csgAB* in Figure 5, is likely from the cellulose component of the EPS matrix that is also controlled by transcriptional regulator CsgD (65). Treatment of the *S. flexneri* 2457T IVLC-induced biofilm with cellulase, which hydrolyzes β-1,4 glycosidic linkages (66), resulted in significant reduction in the IVLC-induced biofilm (**Supplemental Figure S4**). Regardless of the curli disorganization and presence of cellulose, our data demonstrate that the *S. flexneri* 2457T curli are produced and have functional roles in biofilm and epithelial cell adherence.

The pseudogene annotations, particularly for the genes encoding the pores or chaperone-usher components required for assembly of the major structural subunits, warrant future investigations into determining how *S. flexneri* 2457T assembles the adherence structures (Figure 9). If *fimD* is nonfunctional, we hypothesize that homologous genes located in other genomic locations may compensate for a pseudogene in an operon if needed. For example, the *ybgQ, ycbS*, or *yehB* ushers and accompanying chaperone genes may compensate for the truncated expression of *fimD* in the *fim* operon to enable FimA secretion and assembly. This hypothesis is supported by the demonstration of fimbrial promiscuity in biogenesis in *E. coli* in which heterologous structural subunits or secretion systems from different operons are utilized to generate and assemble intact structures (67, 68). Fimbrial promiscuity has also been suggested for *Proteus mirabilis* since soluble Fim14A was detected by MS in the extracellular environment despite an incomplete operon in which the chaperone is absent and the usher is annotated as a pseudogene. *Proteus mirabilis* encodes 17 chaperone-usher fimbrial operons; and therefore, compensation by one of the other operons is hypothesized to enable Fim14A secretion (69). Thus, functional products are likely produced by the other *S. flexneri* 2457T operons, especially for the ushers given the transcriptomic analyses performed and the identification of at least three *fimD* homologs throughout the genome as denoted by the color coding in Figure 3 and **Supplemental Figure S2**.

**Figure 9.**
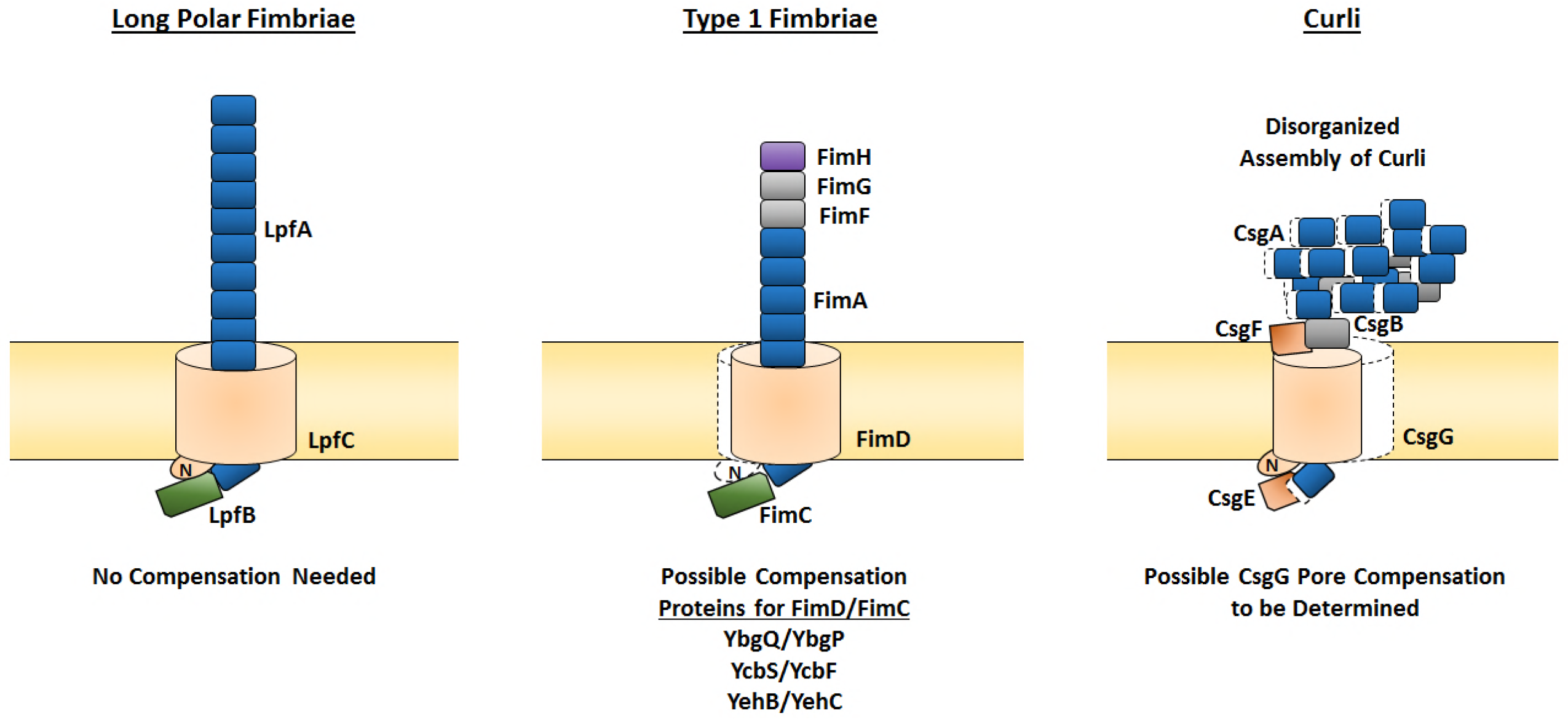
Model of the *Shigella flexneri* adherence factors. Based on the transcriptomic and functional analyses performed, the protein organization for each of the three adherence factors is predicted. First, the major components for the long polar fimbriae are intact and functional. Second, the major and minor structural subunits for type 1 fimbriae are intact to form a full structure. Due to the insertion sequence in *fimD*, we predict either a truncated protein in which the N-terminus is missing (highlighted by the white shadow relative to *E. coli*) or compensation by one of the other FimD homologues listed to enable secretion of the structural subunits. Depending on the truncation, FimC still may be able to interact with FimD or the homologues to serve the appropriate chaperone function. Otherwise, the corresponding FimC homologue would be utilized. Third, due to the N-terminal CsgA truncation and/or the C-terminal CsgG truncation, we predict secretion but not appropriate assembly of CsgA and CsgB for curli. Curli assembly may also be affected by a possible lack of CsgF interaction with a truncated CsgG. Colors for the proteins correspond to the color scheme in Figure 3.

In conclusion, we have demonstrated that *S. flexneri* 2457T produces at least three functional adherence factors for IVLC-induced biofilm formation and adherence to colonic epithelial cells despite the presence of any mutations that would normally inhibit expression. Future investigations, including in-depth analyses defining the mechanism of adherence factor production and secretion in IVLCs as well as studies with other *Shigella* isolates and species, will enhance our understanding of the evolution of this pathogen. Analysis of two clinical *S. flexneri* isolates thus far demonstrated conserved phenotypes (**Supplemental Figure S5**). The pathoadaptive changes that *Shigella* sustained was not the loss of adhesion expression, but rather a precise control of gene expression to enable the production of adhesins only when necessary and in instances that are most beneficial to the pathogen. We agree constitutive expression of these adherence factors would possibly interfere with the pathogenic lifestyle of *Shigella* and impair critical immune evasion tactics. Similar regulation of adhesion genes has been described for other bacterial pathogens such as enterotoxigenic, enterohemorrhagic, and uropathogenic *E. coli* (70–74). Clearly human-adapted pathogens have efficiently evolved to regulate virulence gene expression for efficient colonization and infection tactics in the human host. Our work provides an example of this concept and highlights the importance of utilizing IVLCs to study bacterial pathogens. Finally, this work has profound effects on *Shigella* therapeutic development. The adherence factors provide innovative targets and promise for novel therapies and new strategies to ultimately control and prevent *Shigella* infection.

## Materials and Methods

### Ethics statement

Human sample collection was approved by the institutional review board (IRB) protocol #2015P001908, Massachusetts General Hospital, Boston, MA. Donor tissue was obtained from consenting patients undergoing medically required surgical resections as determined by a licensed physician. All subjects were provided written informed consent.

### Bacterial strains and growth conditions

The list of bacterial strains and plasmids used in this study are presented in Table 3. Bacteria were routinely cultured at 37°C in either Luria broth (LB Lennox) or tryptic soy broth (TSB, which contains additional 2.5 g/L glucose relative to LB), with aeration or in tissue culture treated plates to represent static growth conditions (31, 32). Plating for colony forming units was performed using tryptic soy broth plates with 1.5% agar and 0.025% Congo red (CR, Sigma C6277). Bile salts (Sigma B8756) were used at a concentration of 0.4% w/v. All media were filter sterilized with a 0.22 μM filter following the addition of bile salts and/or glucose. Chloramphenicol was used at 5 μg/ml, kanamycin was used at 50 μg/ml, and ampicillin at 100 μg/ml where indicated.

**Table 3.**
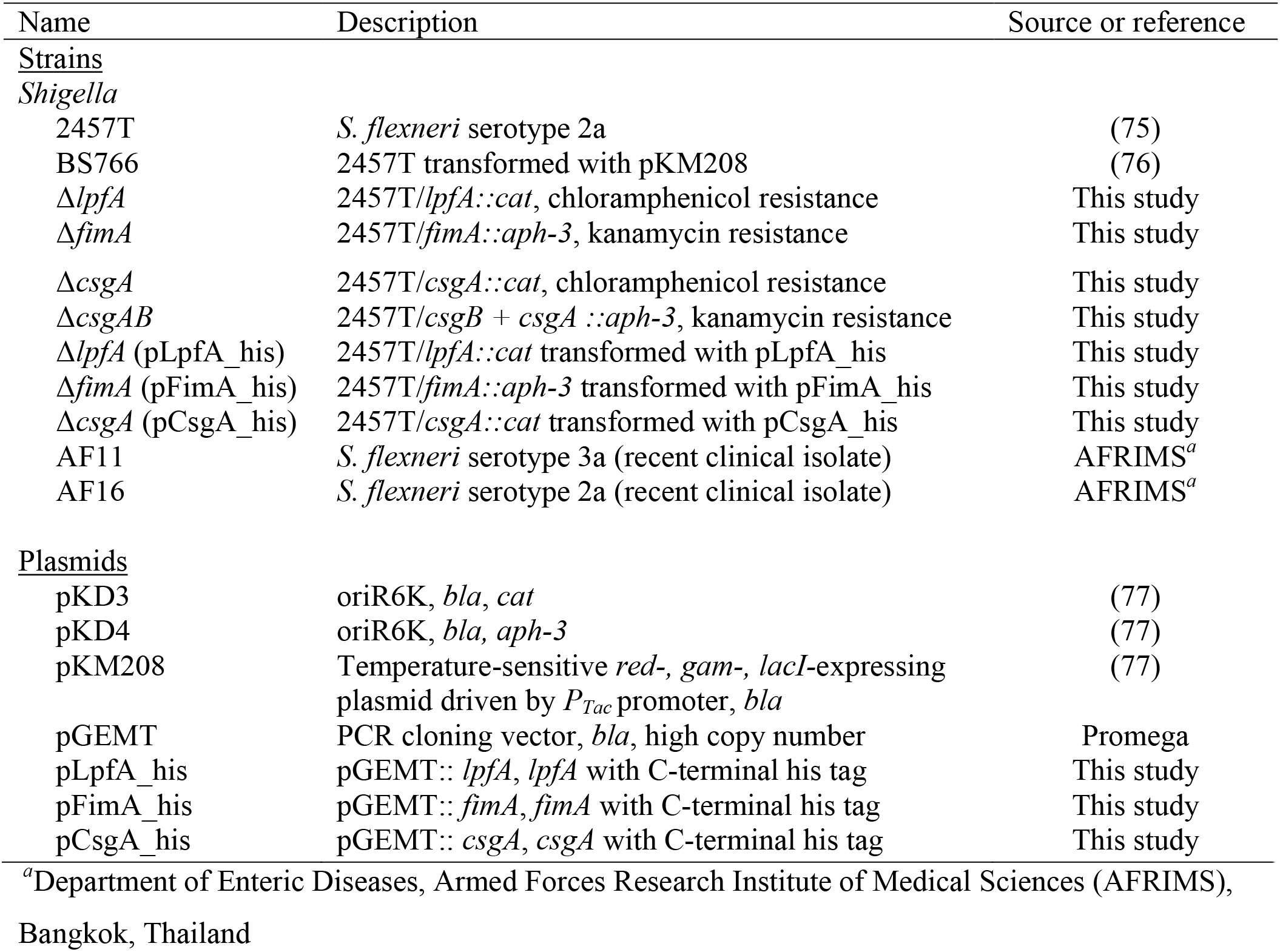
Strains and plasmids used in this study.

| Name | Description | Source or reference |
| --- | --- | --- |
| <u>Strains</u> |  |  |
| *Shigella* |  |  |
| 2457T | *S. flexneri* serotype 2a | (75) |
| BS766 | 2457T transformed with pKM208 | (76) |
| Δ*lpfA* | 2457T/*lpfA::cat*, chloramphenicol resistance | This study |
| Δ*fimA* | 2457T/*fimA::aph-3*, kanamycin resistance | This study |
| $\Delta csgA$ | 2457T/*csgA::cat*, chloramphenicol resistance | This study |
| $\Delta csgAB$ | 2457T/*csgB* + *csgA::aph-3*, kanamycin resistance | This study |
| $\Delta lpfA$ (pLpfA_his) | 2457T/*lpfA::cat* transformed with pLpfA_his | This study |
| $\Delta fimA$ (pFimA_his) | 2457T/*fimA::aph-3* transformed with pFimA_his | This study |
| $\Delta csgA$ (pCsgA_his) | 2457T/*csgA::cat* transformed with pCsgA_his | This study |
| AF11 | *S. flexneri* serotype 3a (recent clinical isolate) | AFRIMS<sup>a</sup> |
| AF16 | *S. flexneri* serotype 2a (recent clinical isolate) | AFRIMS<sup>a</sup> |
| Plasmids |  |  |
| pKD3 | oriR6K, *bla*, *cat* | (77) |
| pKD4 | oriR6K, *bla*, *aph-3* | (77) |
| pKM208 | Temperature-sensitive *red-*, *gam-*, *lacI*-expressing plasmid driven by *P<sub>Tac</sub>* promoter, *bla* | (77) |
| pGEMT | PCR cloning vector, *bla*, high copy number | Promega |
| pLpfA_his | pGEMT:: *lpfA*, *lpfA* with C-terminal his tag | This study |
| pFimA_his | pGEMT:: *fimA*, *fimA* with C-terminal his tag | This study |
| pCsgA_his | pGEMT:: *csgA*, *csgA* with C-terminal his tag | This study |

<sup>a</sup>Department of Enteric Diseases, Armed Forces Research Institute of Medical Sciences (AFRIMS), Bangkok, Thailand

### Biofilm assays

The biofilm assay was performed as previously described (31, 32). Single colonies of each bacterial strain were inoculated into media (LB + 2% glucose or TSB) with bile salts in a single well of a 96-well plate. Plates were incubated at 37°C without shaking. At the indicated time points, wells were gently washed twice with 1X phosphate buffered saline (PBS) either fixed for electron microscopy (see below) or stained with 0.5% crystal violet for 5 minutes. Afterwards, the wells were gently washed five times with sterile dH_2_0 and then set to air dry. Biofilm formation was quantified by adding 95% ethanol to the wells to solubilize the crystal violet stain. After 30 minutes of incubation at room temperature on an orbital shaker at 70 rpm, absorbance at 540 nm (OD_540_) was measured with the plate reader (78). Absorbance readings at OD_600_ were taken to ensure there were no significant differences in growth prior to the washing steps. For experiments in which cellulase (Sigma C1184) was used, 60 units/mL of enzyme were added to wells at the start of the biofilm assay. For complementation analysis, the assays were performed at 4.5 hours to enable appropriate expression of the genes from the pGEMT plasmid. Cellulase and complementation biofilms were subsequently processed as described above. Statistical significance was determined by Student’s t-test (for +/− bile salts comparisons within each strain) or an ANOVA, and a p-value of ≤ 0.05 was considered significant.

### Adherence assays

The isolation and preparation of human intestinal epithelial cells were performed as previously described (34–37, 79, 80). The excess healthy margins of the ascending colon, as verified by a pathologist, were used to obtain the intestinal crypts. The tissue was washed once in cold 1X PBS (ThermoFisher Scientific, MA) and then tissue strips were cut and placed into a dissociation buffer consisting of 1X PBS, penicillin/streptomycin (pen/strep; ThermoFisher Scientific), 1 mM dithiothreitol (DTT; Sigma-Aldrich, MO), and 0.5 mM EDTA (Sigma-Aldrich). Intestinal strips were incubated at 4°C for 30 minutes, then vigorously shaken to promote epithelium dissociation from the basal membrane. This procedure was repeated five times to collect multiple fractions. Subsequently, the fractions containing the intestinal crypts were further processed and plated in Matrigel (Corning, NY) as previously described (34, 79). Intestinal crypt-derived organoids were incubated at 37°C with 5% CO_2_ in media that consisted of a 1:1 ratio of stem cell media and L-WRN (ATCC CRL-3276)-derived conditioned media, in which both media types were prepared as previously described (34, 81). Culture medium was replenished every other day and the organoids were passaged every 7 to 9 days using standard trypsin-based techniques. Approximately 2.0 × 10^6^ cells/ml were re-plated in Matrigel to ensure robust propagation of the organoids (34).

Organoid-derived cell monolayers were generated as previously described (34–37). Single cell suspensions derived from the organoids were plated on polyethylene terephthalate (PET) membrane transwell inserts with a 0.4μm pore size (Corning, NY) at 1.0 × 10^6^ cells/ml and incubated in the 1:1 stem cell/L-WRN media at 37°C with 5% CO_2_. Culture medium was changed every other day until the cultures reached confluence as determined by trans-epithelial electrical resistance (TEER) monitoring and microscopic observation. At 48 hours prior to each experiment, media in the apical chamber were replaced with complete DMEM/F12 plus 5 μM of the γ-secretase inhibitor IX (DAPT; Calbiochem) while the basolateral media were replenished with the 1:1 stem cell/L-WRN media with 10μM Y-27632 (Calbiochem) and 100 to 500 ng/ml of the receptor activator of NF-κB ligand (RANKL; Peprotech). This process was utilized to promote cell differentiation (34, 36), especially for M cells (82). On the day of each experiment, monolayers were washed with 1X PBS, both apical and basolateral media were replaced with DMEM without phenol red, and monolayers were incubated for at least 2 hours before the initiation of the experiment. *S. flexneri* 2457T or the various mutants were subcultured in TSB + bile salts were washed in 1X PBS, resuspended in DMEM without phenol red, applied to the apical surface of the monolayers without centrifugation, and incubated for 3 hours as previously described for polarized T84 cells (30). Afterwards, infected cells were processed for adherence quantification (30) or fixed for electron microscopy (see below). The average percent recovery was calculated for two independent experiments, each with at least two technical duplicates, as [recovered bacterial titer/infecting titer] x 100%. Statistical significance was determined by comparing wild type *S. flexneri* 2457T to each mutant using the Student’s t-test; and, a p-value of ≤ 0. 05 was considered significant.

The HT-29 adherence assay was performed as previously described (31). HT-29 cells (ATCC HTB-38) were seeded in DMEM to establish a semi-confluent monolayer of approximately 75%. For bacterial cultures, single colonies of *S. flexneri* 2457T or the various mutants were inoculated into media (LB) or media with 2% glucose and 0.4% w/v bile salts in tissue culture plates and grown statically at 37°C. Following overnight growth, the bacteria were collected, washed with 1X PBS, standardized to an OD_600_ of 0.35, resuspended in DMEM, and applied to the HT-29 cell monolayers without centrifugation. Cells were incubated at 37°C with 5% CO_2_ for 3 hours. Afterwards, the monolayers were washed five times with 1X PBS and lysed with 1% Triton X-100. Serial dilutions were made to determine the number of cell-associated bacteria. The average percent recovery was calculated for three independent experiments as [recovered bacterial titer/infecting titer] x 100%. Statistical significance was determined by comparing wild type *S. flexneri* 2457T to each mutant using the Student’s t-test; and, a p-value of ≤ 0.05 was considered significant.

### Electron microscopy analyses

For the biofilm culture analysis, single colonies of *S. flexneri* 2457T or the various mutants were added to tissue culture-treated plates containing LB media or LB media supplemented with a final concentration of 2% w/v glucose and/or 0.4% w/v bile salts. Cultures were grown statically overnight at 37°C. On the following day, samples were collected and prepared for transmission electron microscopy (TEM) imaging by fixing in 2.5% glutaraldehyde and staining with uranyl acetate (83). Samples were mounted on Formvar/Carbon 100 Mesh grids (Electron Microscopy Services) and imaged with a JEOL transmission electron microscope. For scanning electron microscopy (SEM) analysis of the HIODEM adherence assay, samples were fixed in 0.5X Karnovsky fixative and subsequently stored in 1X PBS. All sample processing occurred at the Massachusetts Eye and Ear Infirmary core facility. All SEM imaging was performed at the Harvard University Center for Nanoscale Systems (CNS) using a FESEM Supra55VP microscope. The SEM images were pseudo-colored according to protocols listed at http://www.nuance.northwestern.edu/docs/epic-pdf/Basic_Photoshop_for_Electron_Microscopy_06-2015.pdf.

For TEM analysis of isolated adherence factors, wild type and mutant strains were cultured statically in LB plus 2% glucose and 0.4% w/v bile salts, and an ammonium sulfate precipitation was performed (47). Briefly, samples were collected and pelleted by centrifugation at 4000 rpm for 10 minutes. The bacterial pellet was resuspended in 1X PBS and heated at 65°C for 30 minutes and subsequently centrifuged at 4000 rpm for 10 minutes. The supernatants were transferred to a new tube and precipitated by mixing the samples with 40% ammonium sulfate on an end-over-end mixer for 10 minutes at room temperature. Afterwards, the samples were dialyzed in 1X PBS using 3.5 MWCO dialysis cassettes for 1h at RT on a 50 RPM rotating shaker. The 1X PBS was then changed and the cassettes were transferred to 4°C for overnight dialysis. The dialyzed fraction was collected and stored at -20°C. A fraction of each sample was fixed and processed for TEM analysis.

### RNA isolation

RNA was isolated from bacterial cultures as previously described (84) with Qiagen’s RNeasy kits. DNA was digested with Turbo DNase (Invitrogen), and concentrations of total RNA were determined using a NanoDrop ND-1000 spectrophotometer. The cDNA was synthesized from total RNA using Superscript III First Strand Synthesis kit (Invitrogen) or RevertAid cDNA first strand synthesis kit (Thermo Scientific) according to manufacturers’ protocols. All RNA was first confirmed to be free of DNA contamination by performing separate cDNA synthesis reactions with and without reverse transcriptase, followed by PCR amplification of the house-keeping gene *rpoA* as described previously (31).

### RNA sequencing (RNA-seq) analysis

The data generated from the RNA-seq analysis of *S. flexneri* 2457T RNA isolated from broth cultures were performed in our previous study (31). Duplicate cultures were grown either statically or shaking aeration in TSB or TSB supplemented with 0.4% bile salts as previously described. The RNA-seq trace read data were generated using the Integrative Genomics Viewer (IGV) software version 2.3.67 (85, 86). Each RNA-seq data set was loaded into the IGV software and the traces were normalized to the *S. flexneri* 2457T *rpoA* gene on the autoscale setting. The zoomed-in traces for two genes provided in Supplemental Figure 1 represents a 10-fold magnification in the scale setting. Genes of interest were searched using the publicly available *S. flexneri* 2457T genome (GenBank Accession number AE014073.1) and *S. flexneri* 2a strain 301 virulence plasmid annotations (GenBank Accession number AF386526.1).

### Reverse-transcription PCR (RT-PCR) analysis

For non-quantitative RT-PCR analysis, cDNA was synthesized from total RNA isolated from broth cultures using the RevertAid cDNA first strand synthesis kit (Thermo Scientific) according to manufacturer’s protocol. All RNA was first confirmed to be free of DNA contamination as described above. The various PCR reactions were performed using the 2X Taq-Pro Complete PCR mix (Denville Scientific). All primer sets were validated and tested for proper DNA amplification prior to the experiment (data not shown). The annealing temperatures were adjusted accordingly for each primer set, and the extension time was adjusted for the size of each product. The products of the reactions were visualized by gel electrophoresis on 1% agarose gels stained with ethidium bromide on a Syngene GelDoc system. The molecular weight markers used in the analysis included GeneRuler, 1 kb plus, and 100 bp plus (Thermo Fisher Scientific). For quantitative RT-PCR analysis (qRT-PCR), biologically independent RNA samples were isolated and ensured DNA-free as described above. Analysis by qRT-PCR was performed as previously described (84), and all data were normalized to levels of *rpoA* and analyzed using the comparative cycle threshold (ΔC_T_) method (87). The expression levels of the target genes under the various conditions were compared using the relative quantification method (87). Real-time data are expressed as the relative changes in expression levels compared with the media without glucose and/or bile salts. Statistical significance was determined using the Student’s t-test to compare each gene expression in control versus treatment media, and a p-value of ≤ 0.05 was considered significant. Due to the significant number of primers used in this analysis, primer sequences are available upon request.

### Mutant construction

The mutants used in this study were constructed using the λ red linear recombination method as previously described (76). Briefly, PCR was used to amplify a chloramphenicol resistance cassette gene (*cat*) from pKD3 or the kanamycin resistance gene cassette *(aph-3)* from pKD4 (Table 1) with 5’ and 3’ overhangs identical to the 5’ and 3’ regions of each gene of interest. Antibiotic resistant recombinants were identified and selected on chloramphenicol or kanamycin plates, and subsequently screened via PCR using confirmation primers that annealed to unique regions up and downstream of each gene to detect the size difference due to the insertion of the antibiotic cassette. Primer sequences for the mutant construction and confirmation are also available upon request.

### Plasmid construction

The plasmids encoding the histidine-tagged LpfA, FimA, and CsgA were constructed as previously described (30). Briefly, each gene and respective native promoter regions were amplified by PCR with high fidelity Taq polymerase (Invitrogen) from wild type 2457T. For FimA, a 6X his tag was added to the C-terminus followed by a stop codon. For LpfA and CsgA, a glycine linker sequence was added upstream of the 6X his tag. The PCR products were ligated into pGEMT and the plasmids were subsequently transformed into the appropriate adherence mutant. Selection for positive transformants occurred on tryptic soy broth plates containing 1.5% agar, 0.025% Congo red, and 100 μg/ml ampicillin. Sequencing was performed to ensure no mutations were introduced during the cloning process. All primers used for the plasmid constructions and sequencing verification will also be made available upon request.

### Congo red binding assay for curli detection

Samples for the Congo red binding assay were collected by gentle scraping from the biofilm and processed for ammonium sulfate precipitation as detailed above, and placed on a clean, dry glass slide. The specimens were air-dried, subsequently stained with alkaline Congo red solution (Sigma HT603), and incubated at room temperature for approximately 10 minutes. The smears were rinsed in water until excess stain was drained and the slides were observed under polarized light for apple green birefringence (49, 50). Samples were imaged with a Nikon Ci-E microscope with an attached camera.

### Mass spectrometry analysis

*Shigella flexneri* 2457T was cultured in TSB + 0.4% w/v bile salts as described above for the biofilm assay. Following o/n incubation, culture supernatants were collected and concentrated by trichloroacetic acid (TCA) precipitation. The protein pellet was stored at -20°C until analyzed. For mass spectrometry (MS) analysis, first, intact mass analysis was performed by reconstituting the lyophilized sample in 0.1% trifluoroacetic acid. UPLC-QToF MS was performed to detect the masses of intact molecules present in the mixture. Samples were analyzed using reversed-phase liquid chromatography (RPLC) and a Xevo G2-S Q-TOF (Waters Corp, Milford, MA). Liquid chromatography was performed at 0.200 mL/min using an H-Class Acquity ultra-high pressure liquid chromatography system (UPLC; Waters Corp, Milford, MA) on a BEH300-C4 column (2.1 mm x 150 mm, pore size of 1.7 μm; Waters Corp, Milford, MA). Buffer A consisted of 0.1 % formic acid (vol/vol) in UPLC grade water and buffer B consisted of 0.1 % formic acid (vol/vol) in 100 % UPLC grade acetonitrile. In all analyses, a gradient separation was performed as follows: 0 min 90% A, 5 min 90% A, 80 min 10% A, 90 min 10% A, 91 min 90% A, 100 min 90% A. After RPLC, samples were introduced via an electrospray ion source inline with the Xevo G2-S Q-TOF. External calibration of *m/z* scale was performed using sodium cesium iodide. The Q-TOF parameters were run in sensitivity mode, scanning m/z 400-4000, 3.00 kV capillary voltage, 40 V cone voltage, 150°C source temperature, 350°C desolvation temperature, and 800 L/h desolvation gas. MS data were collected at a scan speed of 1.0 s. LC solvents were UPLC grade and all other chemicals were of analytical grade. Intact masses were calculated using the Waters UNIFI software package and deconvolved using the MaxEnt algorithm.

For peptide analysis, samples were digested with trypsin at 37°C for 1.5 hours and the resulting peptides were subsequently extracted for analysis. UPLC-QToF MS/MS was performed to detect the masses of digested peptides and the respective fragments. Samples were analyzed using RPLC as described on a BEH300-C18 column (2.1 mm x 150 mm, pore size of 1.7 μm; Waters Corp, Milford, MA) using the same Buffer A and Buffer B compositions. In all analyses, a gradient separation was performed as follows: 0 min 95% A, 2 min 95% A, 55 min 40% A, 64 min 10% A, 74 min 10% A, 75 min 95% A, 90 min 95% A. After RPLC, samples were introduced via an electrospray ion source inline with the Xevo G2-S Q-TOF. External calibration of *m/z* scale was performed using sodium cesium iodide. The Q-TOF parameters were run in resolution mode, scanning m/z 50-2000, 3.00 kV capillary voltage, 30 V cone voltage, 130 °C source temperature, 250 °C desolvation temperature, and 800 L/h desolvation gas. MS/MS data were collected at a scan speed of 0.1 s. LC solvents were UPLC grade and all other chemicals were of analytical grade. Peptide fingerprinting was completed through the Waters UNIFI software package. Parameters were set to restrict matches only to those peptide fragments where the primary ion exhibited > +1 charge and at least 1 daughter ion was detected confirming the presence of each particular peptide. Any peptide maps with less than 10% coverage were excluded from the analysis.

### Immunoprecipitation analysis

Each strain harboring the his-tagged constructs (Table 3) were grown in static overnight biofilm cultures as described above. For plasmid maintenance, ampicillin was added. Culture supernatants were subsequently collected, filtered sterilized, and TCA precipitated. The total protein pellets were resuspended in 1 mL NP-40 with protease inhibitor cocktail (Roche Diagnostics GmbH). Samples were pre-cleared using Protein A/G plus agarose beads (Pierce) followed by immunoprecipitation with a mouse anti-his affinity resin (Genescript) or a negative control mouse IgG antibody (Santa Cruz). Samples were incubated overnight at 4°C with rotation. On the following day, protein A/G plus agarose beads were added to the negative IgG control samples and incubated for 1 hour at 4°C with rotation. Afterwards, the beads or resin samples were pelleted, washed six times, and boiled in acidified Laemelli lysis buffer as previously described for adherence proteins(88). After boiling, samples were neutralized with basic Laemelli lysis buffer. Samples were run on a 4-20% SDS-PAGE gel (Biorad) and Western blot analysis was performed as previously described (30) using a primary anti-his antibody (Qiagen) and a secondary Alexa Fluor 700 goat anti-mouse antibody. Western blots were scanned using the Odyssey infrared detection system (Li-Cor).

## Acknowledgements

We would like to thank Dr. Stefania Senger, Dr. Alessio Fasano, and Ms. Laura Ingano for use and assistance with the human intestinal organoid-derived epithelial monolayer model. We would also like to thank Dr. Brett Swierczewski, Walter Reed Army Institute of Research, for the clinical isolates of *S. flexneri* used in this study. This work was supported by the National Institute of Allergy and Infectious Diseases grants K22-AI104755 (CSF), 5T32-AI095190-04 (JRS), and U19-AI110820 (DAR). Funding for BJDK and the Mucosal Immunology and Biology Summer Center’s Internship Program is provided by the National Institute of Diabetes and Digestive and Kidney Diseases grant R25 DK103579. Funding for the human intestinal organoid-derived epithelial monolayer model is supported by the National Institute of Allergy and Infectious Diseases grants U19-AI082655. The TEM core is supported by National Institute of Neurological Disorders and Stroke P30-NS045776. Support for the Philly Dake Electron Microscope Facility was provided by the National Institutes of Health grant 1S10RR023594S10 and by funds from the Dake Family Foundation. The funders had no role in study design, data collection and analysis, decision to publish, or preparation of the manuscript. The content is solely the responsibility of the authors and does not necessarily represent the official views of the funders.

## Supporting Information

**Supplemental Figure S1. Putative adherence factor production in TSB + bile salts**. Wild type cultures of 2457T were grown overnight with static growth in TSB media supplemented with 0.4% w/v bile salts. TEM analysis revealed similar structures compared to bacteria grown in LB supplemented with 2% glucose and 0.4% bile salts. Image viewed at 50,000 X magnification (500 nm scale bar).

**Supplemental Figure S2. Additional adherence gene cluster analysis**. Analysis was performed as described in the Materials and Methods as well as in the Figure 2 legend. A 10-fold enhanced view of the trace reads are provided for genes S3342 and *sfmA* into *sfmC* to demonstrate the presence of RNA-seq reads.

**Supplemental Figure S3. The Congo red binding assay**. The assay was performed to confirm the presence of curli due to the ability of the Congo red dye to bind amyloid fibers and produce a birefringence signal under polarized light. The apple-green color indicates Congo red fluorescence that occurs when amyloid fibers are present. Wild type 2457T produced a positive signal while a significant reduction in signal was detected for the *ΔcsgA* mutant. The *ΔlpfA* mutant was analyzed as a mutation control; and as seen, the mutation did not affect the birefringence signal. Images are representative of at least two biological independent experiments.

**Supplemental Figures S4. Treatment of the IVLC-induced biofilm with cellulase**. The biofilm formation analysis was performed +/− cellulase to analyze the contribution of cellulose. A significant reduction in biofilm formation was detected in the presence of cellulase. All data represent three the average OD_540_ from three biological independent experiments in which each experiment had technical triplicates. The p-value for the comparison was <0.001 (***).

**Supplemental Figure S5. Analysis of clinical isolates of *Shigella flexneri*.** Both TEM (top) and biofilm formation (bottom) analyses were performed with clinical isolates of *S. flexneri* serotype 3a (strain AF11) and *S. flexneri* serotype 2a (strain AF16) to confirm the results of the study. For the TEM, the image on the left for AF11 is 50,000X magnification while the image on the right for AF16 is 25,000X magnification. For the biofilm assay, plotted is the average OD_540_ for the strains were grown in either TSB (light blue bars) or TSB supplemented with bile salts (dark blue bars). All data represent three biological independent experiments in which each experiment had technical triplicates. The p-values for TSB versus TSB + bile salts for each strain were <0.001 (***).

