## Supplementary figures and images for "*Shigella flexneri* adherence factor expression in *in vivo*-like conditions"

### Supplemental Figure 1

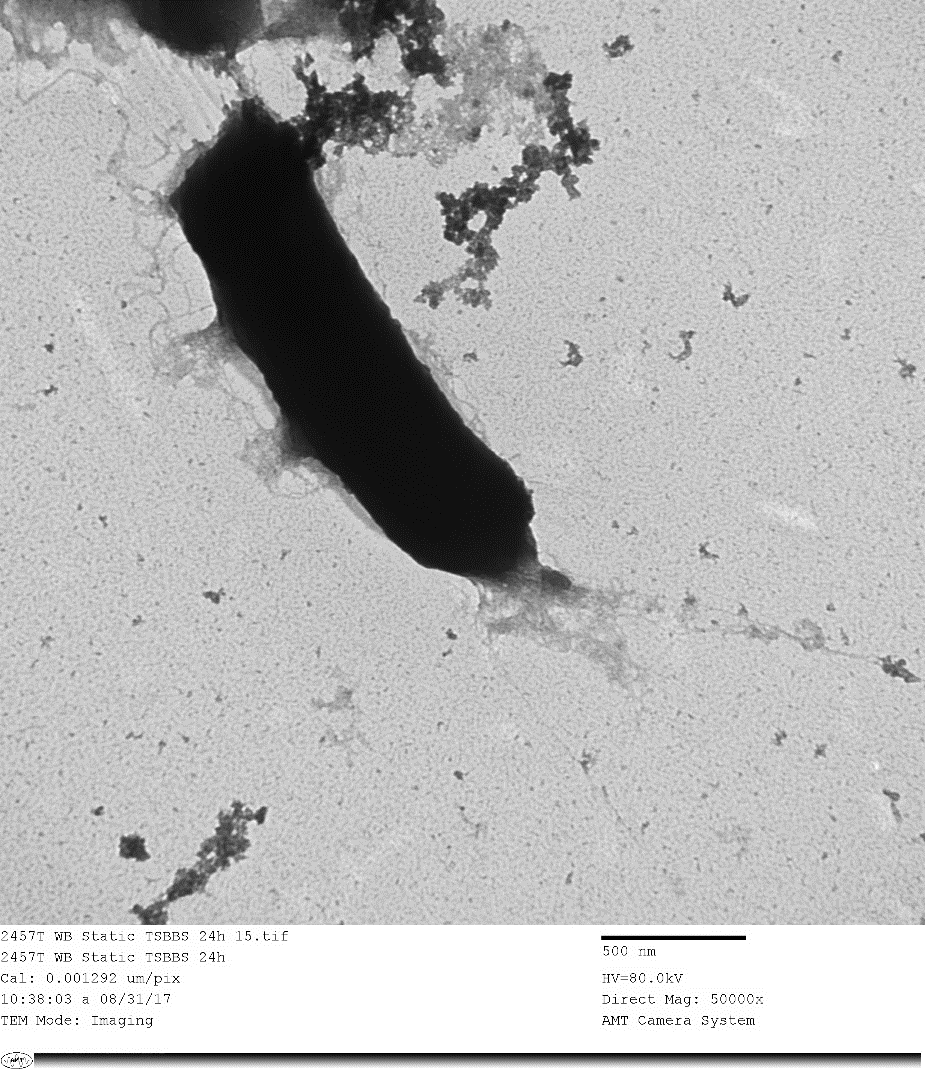
**Supplemental Figure S1**

### Supplemental Figure 2

**Supplemental Figure S2**

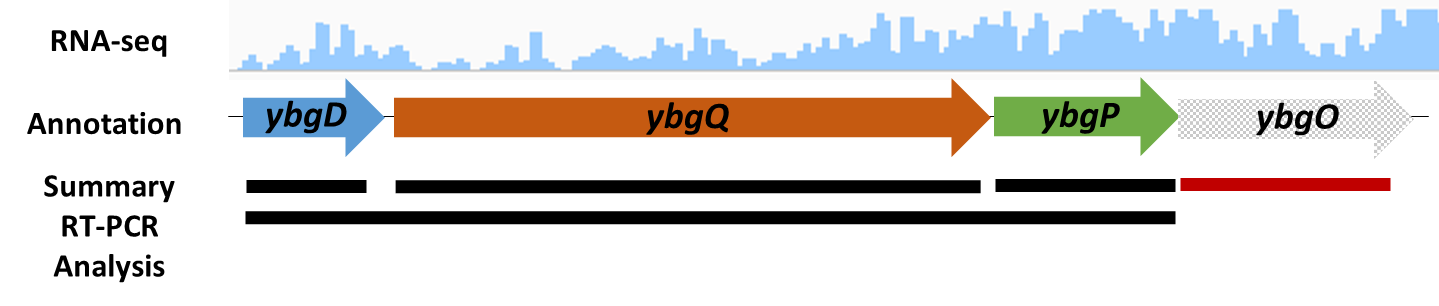

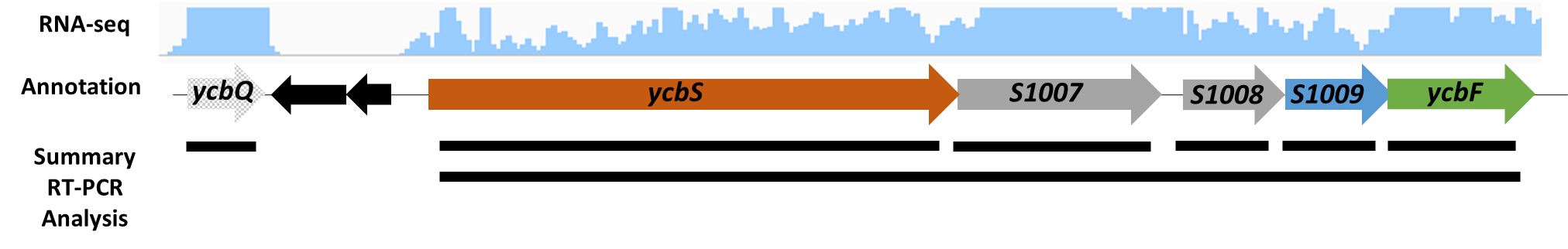

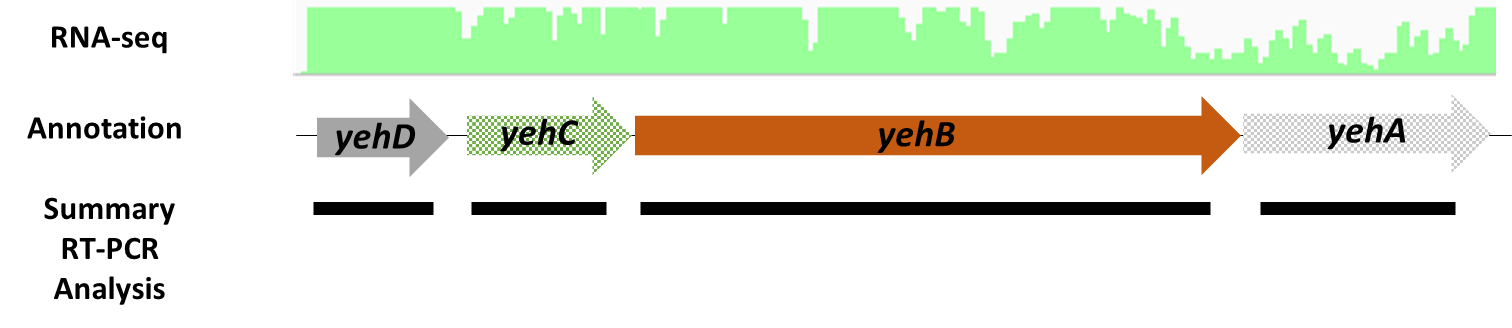

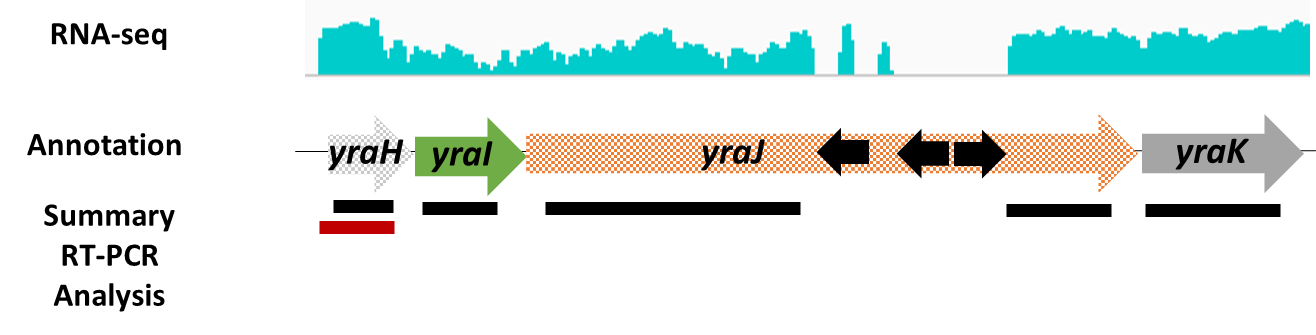

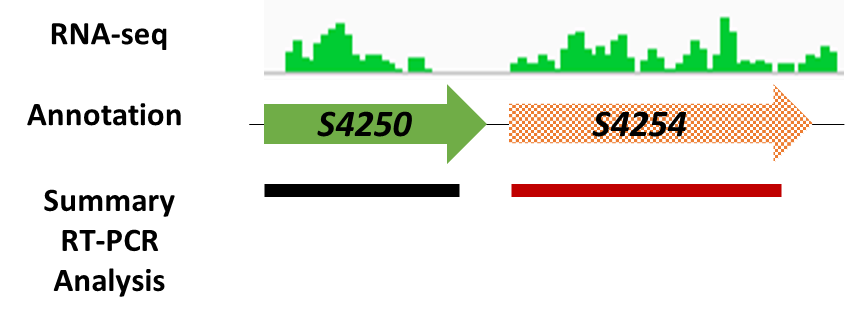

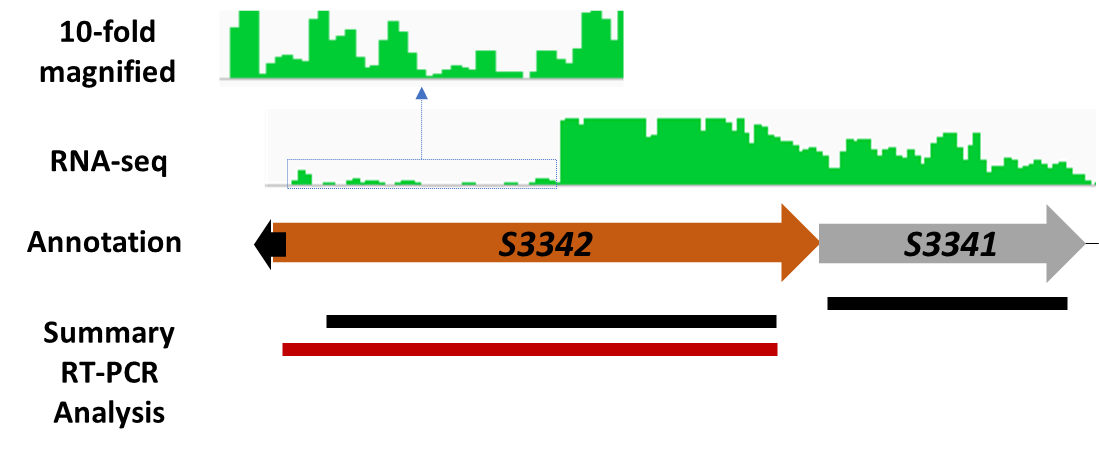

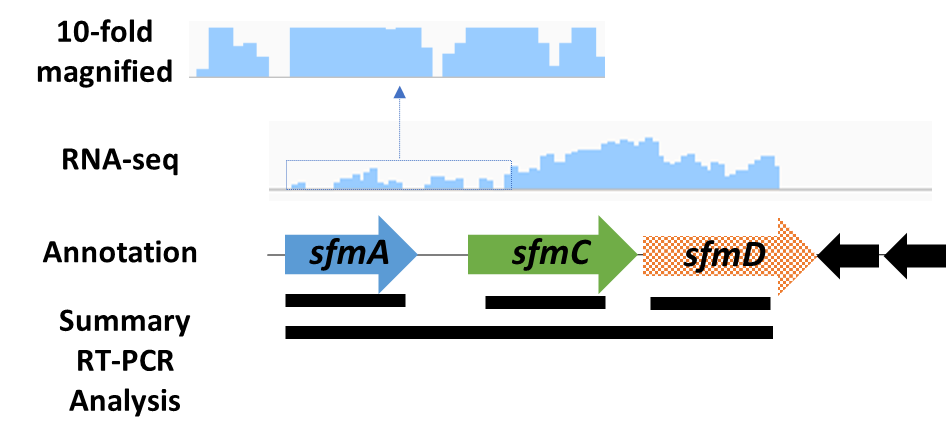

### Supplemental Figure 3

**Supplemental Figure S3**

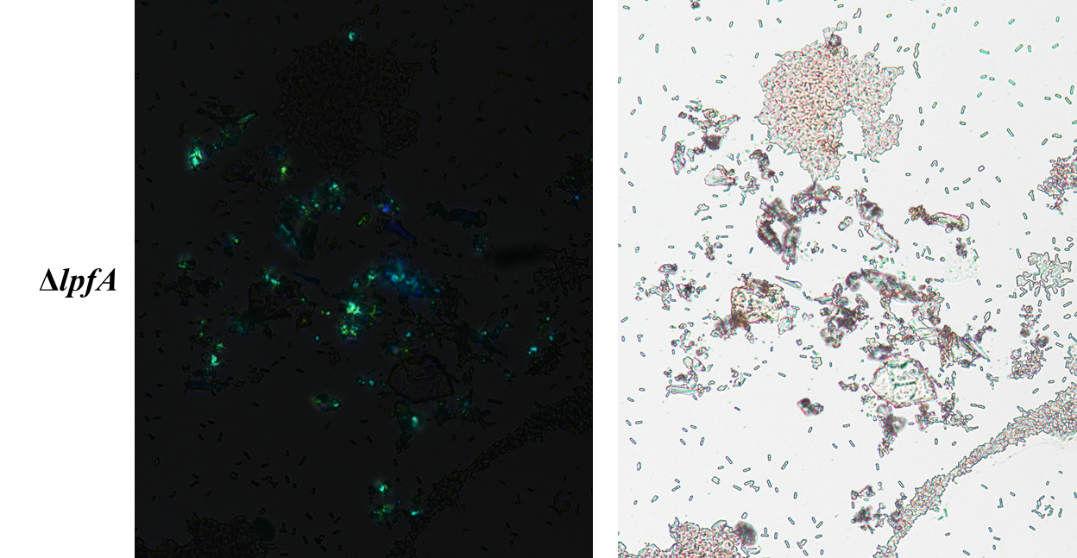

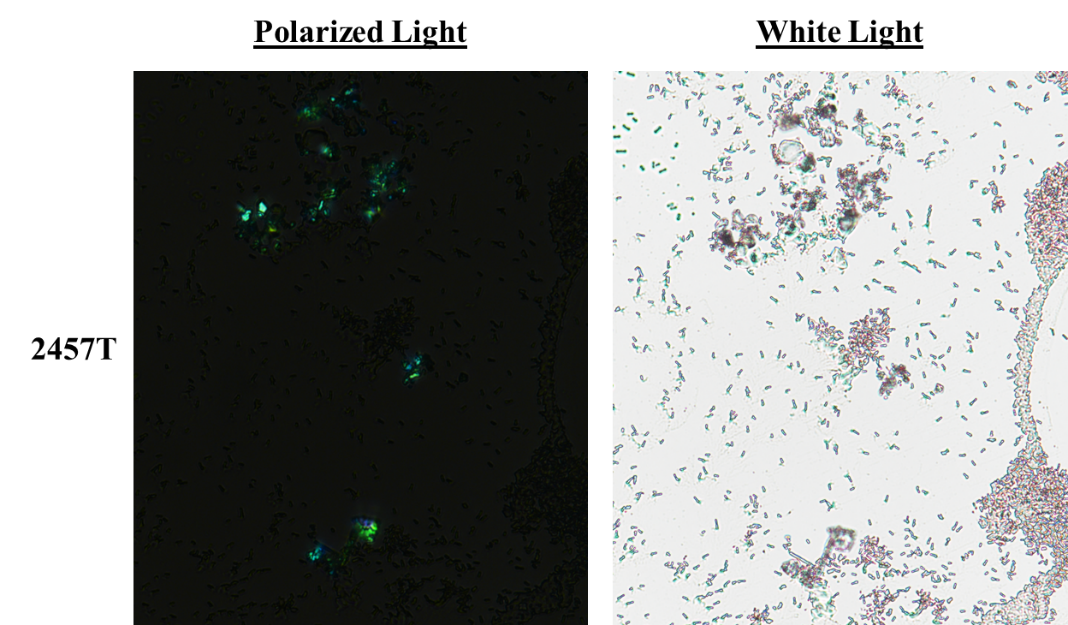

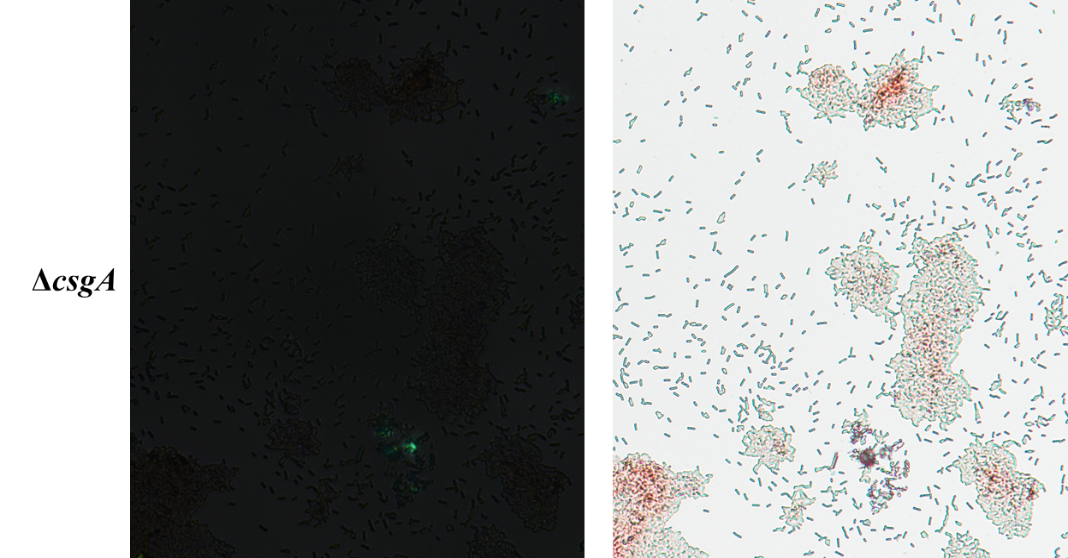

### Supplemental Figure 4

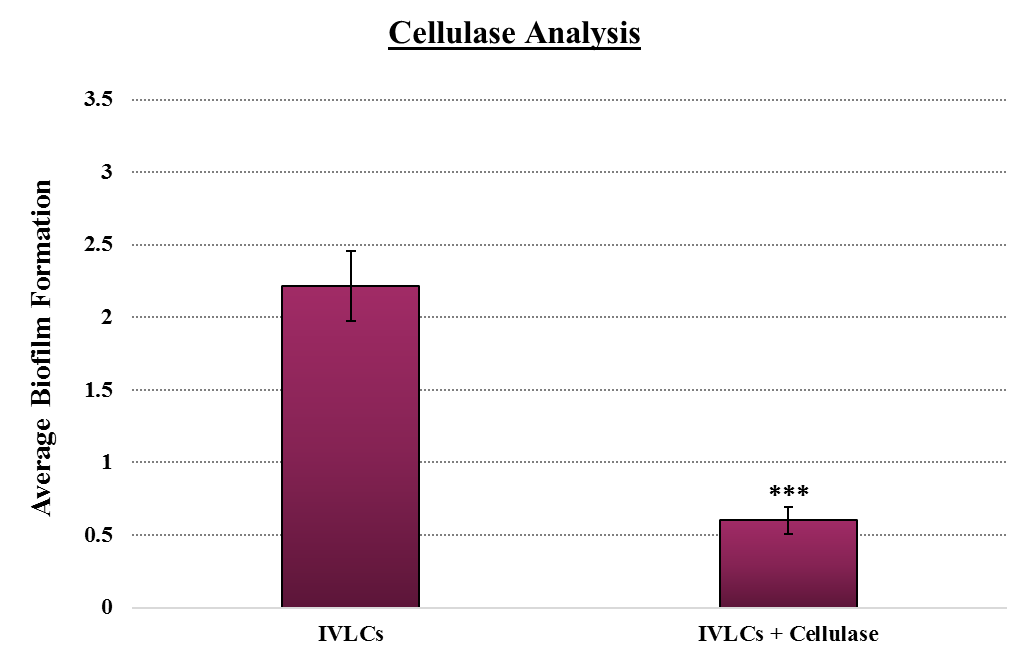
**Supplemental Figure S4**

### Supplemental Figure 5

**Supplemental Figure S5**

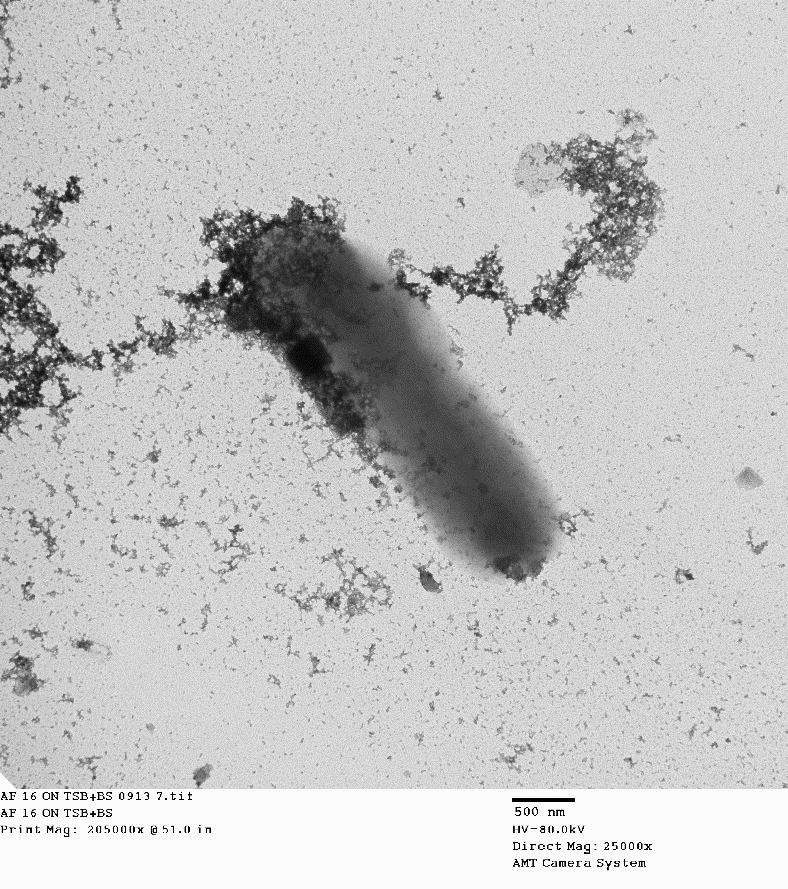

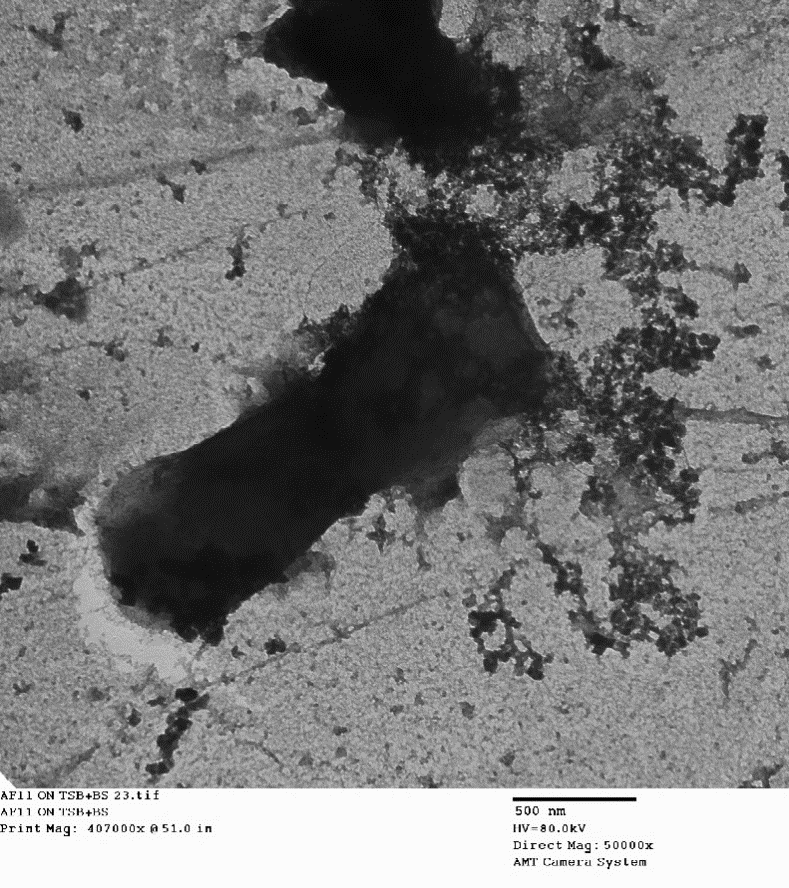

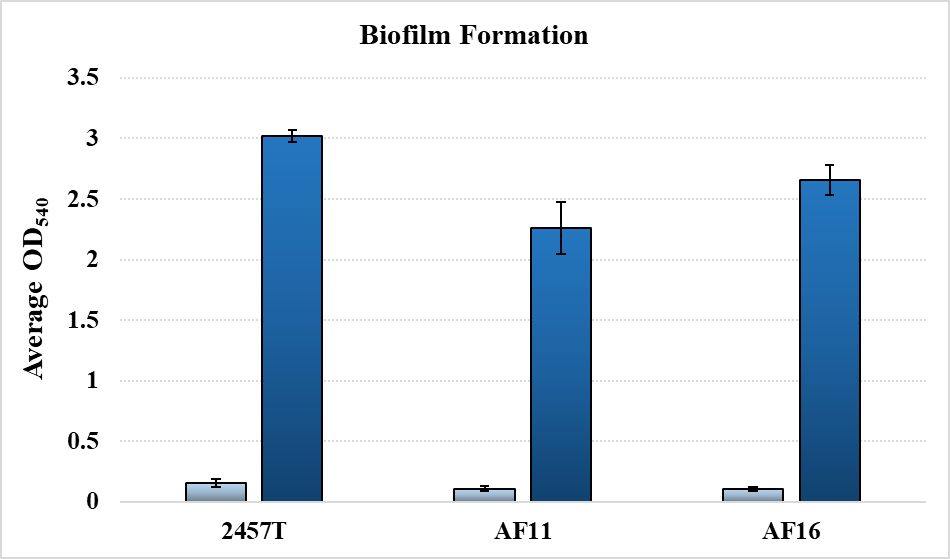

**AF16: *Shigella flexneri* serotype 2a**

**AF11: *Shigella flexneri* serotype 3a**

***

***

***
